# *Mtor*/*Rptor* Function Globally Prevents Cortical Microcephaly and Cell-autonomously Promotes Postnatal Neuron Survival in Cell Type Specific Manner

**DOI:** 10.64898/2026.05.01.722172

**Authors:** Ana Villalba, Robert Beattie, Florian M. Pauler, Carmen Streicher, Osvaldo A. Miranda, Thomas Krausgruber, Martin Senekowitsch, Matthias Farlik, Christoph Bock, Thomas Rülicke, Simon Hippenmeyer

## Abstract

The generation of faithful cell-type diversity and correct projection neuron numbers is essential for cerebral cortex development. Corticogenesis is however susceptible to genetic interference of critical signaling pathways, including mutations in Mtor/Rptor that lead to microcephaly. How the loss of Rptor/mTORC1 function affects cortical developmental programs, at single cell level, is still unknown. Here, we utilized Mosaic Analysis with Double Markers (MADM) technology to probe Rptor gene function upon sparse single cell- or global tissue-wide ablation. We found that tissue-wide effects drive the etiology of cortical microcephaly upon loss of Rptor, rather than deficits in projection neuron genesis. Conversely, Rptor function is cell-autonomously required for postnatal projection neuron survival in a highly cell-type-specific manner. Collectively, our results suggest that the fine balance of precise cell-type-specific cell-autonomous Rptor/mTORC1 function in concert with non-cell-autonomous tissue-wide effects is essential for the development of a properly-sized cerebral cortex with accurate projection neuron diversity.

## INTRODUCTION

The cerebral cortex is organized into six cytoarchitectonic layers and composed of a vast number of neuronal and glial cell types, assembling into cortical circuits that account for major cognitive abilities (Molnar et al. 2019; Hanganu-Opatz et al. 2021; Namba and Huttner 2024; Nowakowski et al. 2025). Based on a variety of criteria including laminar position, morphology, physiological parameters and gene expression pattern, remarkable heterogeneity among cortical projection neuron cell types has been described (Lodato and Arlotta 2015; Zeng 2022; Di Bella et al. 2024; Steyert et al. 2025). The cellular and molecular mechanisms generating appropriate cortical cell-type diversity are however not well understood, albeit at the quantitative level, tightly orchestrated developmental programs ensure the generation of a cerebral cortex of correct size. Importantly, these programs, controlling the faithful generation and maturation of postmitotic neurons by neural stem cells (NSCs), need to be executed precisely. Impairments in cortical neurogenesis may lead to dramatic morphological and scale alterations in the developing cerebral cortex leading to microcephaly (smaller brain) or macroencephaly (bigger brain) (Barkovich et al. 2012; Pirozzi et al. 2018; Asif et al. 2023). These cortical malformations are thought to reflect some aspects of the underlying etiology for neurodevelopmental disorders such as autism, intellectual disability and epilepsy (Sullivan and Geschwind 2019; Klingler et al. 2021; Bizzotto and Walsh 2022).

During development, the cortical cell wall emerges from neuroepithelial stem cells (NESCs) which initially amplify their pool but then transform into radial glial progenitor (RGP) cells (Kriegstein and Alvarez-Buylla 2009; Taverna et al. 2014; Villalba et al. 2021). RGPs are the major neural progenitors and their proliferation dynamics along temporal lineage progression determine the final number of projection neurons in the mature cortex (Taverna et al. 2014; Andrews et al. 2022; Casingal et al. 2022; Mosti et al. 2025). Although we lack a mechanistic understanding of temporal RGP lineage progression at single cell level, recent clonal analysis using Mosaic Analysis with Double Markers [MADM, (Zong et al. 2005; Contreras et al. 2021)] revealed an inaugural quantitative framework of RGP-mediated cortical neurogenesis (Gao et al. 2014; Llorca et al. 2019), with temporally-predictable RGP transition nodes and projection neuron output (Lin et al. 2021; Llorca and Marin 2021; Hippenmeyer 2023). Yet, the cellular and molecular mechanisms regulating stereotyped RGP lineage progression and neurogenic potential at the single RGP cell level remain mostly unknown.

Here, we focus here on the role of mTOR (mammalian Target of Rapamycin) signaling. mTOR is a serine/threonine protein kinase that assembles into two distinct multi-protein signaling complexes – mTORC1 and mTORC2 (Loewith et al. 2002) – and regulates a wide range of metabolic and cellular processes, including cell growth (Liu and Sabatini 2020; Battaglioni et al. 2022). The mTORC1 complex, and downstream signaling via RPTOR [regulatory associated protein of MTOR complex 1; (Hara et al. 2002; Kim et al. 2002)], has been critically associated with the regulation of brain size during development, albeit at the macroscopic whole tissue level (Crino 2011; Magri and Galli 2013; Lipton and Sahin 2014; Pirozzi et al. 2018; Girodengo et al. 2022). In effect, mutations in human that lead to overactivation of mTOR signaling have been shown to result in aberrant larger brain size and cortical malformation focal cortical dysplasia (FCD) (D’Gama et al. 2017; Blumcke et al. 2021; Bizzotto and Walsh 2022; Girodengo et al. 2022; Gerasimenko et al. 2023). Conversely, conditional deletion of *Mtor* (Ka et al. 2014) or *Rptor* (Cloetta et al. 2013) in mice results in severe microcephaly. Thus, mTORC1 appears to fulfill critical functions during development across different species to ensure the buildup of a correctly-sized cerebral cortex. However, the functional requirement for mTOR signaling in RGP-mediated neurogenesis, the generation of cortical projection neuron diversity, and cell-type specific roles during cortical development at the single cell level, is not known. To this end, we capitalized upon the high-resolution MADM technology, enabling quantitative lineage tracing and assessment of RGP-derived projection neuron output in combination with genetic manipulation of *Rptor*/mTORC1 at true single cell level.

## RESULTS

### Clonal analysis reveals that *Rptor* is not cell-autonomously required for RGP-mediated neocortical neurogenesis

Brain-wide deletion of *Rptor* led to microcephaly (Cloetta et al. 2013) but how the loss of *Rptor* affects RGP lineage progression and/or projection neuron output at the individual progenitor cell level remains unclear (Figure 1A-1B). To assess the consequences of *Rptor* loss-of-function with single RGP resolution we conducted MADM-based clonal analysis (Gao et al. 2014; Beattie et al. 2020), enabling precise quantitative assessment of RGP-mediated neurogenesis. We conducted two MADM assays, both in combination with tamoxifen (TM)-inducible *Emx1*-CreER driver (Kessaris et al. 2006), targeting neocortical RGPs.

**Figure 1.**
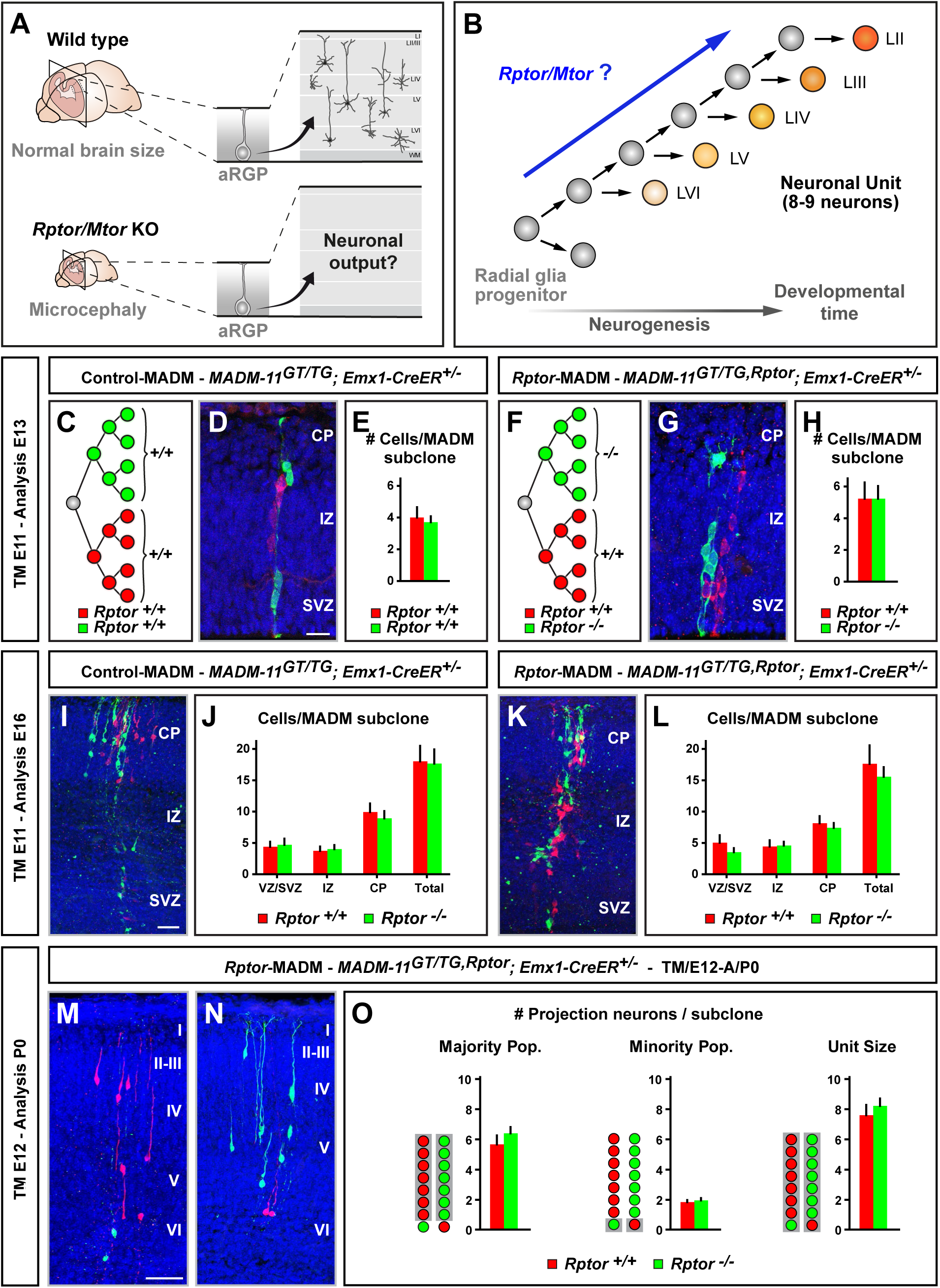
MADM-based clonal analysis reveals that *Rptor* is not cell-autonomously required for cortical RGP-mediated neurogenesis. (A) Illustration of a wild-type and a *Rptor/Mtor* cKO microcephalic mouse brain with possible implications at the level of RGP-mediated projection neuron output. (B) Quantitative framework of RGP lineage progression with a putative functional role of *Rptor/Mtor* pathway. (C-L) MADM clonal analysis in Control-MADM (C-E, I-J; *MADM-11^GT/TG^*;*Emx1-CreER^+/-^*) and *Rptor*-MADM (F-H, K-L; *MADM-11^GT/TG,Rptor^*;*Emx1-CreER^+/-^*) with TM injection at E11 and analysis at E13 (C-H) and E16 (I-L). (C and F) Schematic MADM paradigm for Control-MADM (C) and *Rptor*-MADM (F). Upon the MADM event, two differentially-labeled lineages emerge from a dividing RGP: one red tdT^+^ control lineage in both Control-MADM and *Rptor*-MADM, and one green GFP^+^ control (Control-MADM) or *Rptor*^-/-^ mutant (*Rptor*-MADM) lineage. (D, G, I and K) Representative images of E11-E13 (D and G), and E11-E16 (I and K) MADM clones. (E, H, J and L) Quantification of total clone size at E13 (E and H) and E16 (J and L), and relative distribution in VZ/SVZ, IZ and CP at E16 (J and L). (M-O) MADM clonal analysis in *Rptor*-MADM (*MADM-11^GT/TG,Rptor^*;*Emx1-CreER^+/-^*) with TM injection at E12 and analysis at P0 to determine the clonal projection neuron unit output in asymmetric neurogenic clones. (M and N) 3D reconstruction images of representative asymmetric G2-X MADM clones with majority population in red (M) or green (N) in *MADM-11^GT/TG,Rptor^*;*Emx1-CreER^+/-^*. Note that control cells are labeled in red (tdT^+^) color and *Rptor*^-/-^ cells in green (GFP^+^) color, regardless of whether in majority or minority populations. (O) Quantification of the size of the majority population, minority population and the unitary size in asymmetric neurogenic MADM clones. Nuclei were stained using DAPI (blue). CP: Cortical Plate. IZ: Intermediate zone. SVZ: Subventricular zone. Bars and error bars represent mean ± SEM. No comparison was significant as determined by using unpaired t test (E, H and O) or two-way ANOVA with Šídák’s multiple comparisons (J and L). Scale bar, 20 µm (D and G), 50 µm (I and K), 500 µm (M and N).

In the first experiment (Figure 1C-1L), we analyzed RGPs in their symmetric proliferation mode, whereby the two daughter cells (and emerging subclones) of a dividing RGP will be labeled in red *or* green color, respectively (Figure 1C and 1F). We first generated MADM clones with wild-type genotype which we term Control-MADM [*MADM-11^GT/TG^*; *Emx1-CreER^+/-^* (Figure 1C-E)] to establish a baseline, and injected TM at embryonic day 11 (E11) with most RGPs in symmetric proliferation mode. We quantitatively assessed cell numbers in red and green colors (both wild-type) at E13 (Figure 1D-1E) and E16 (Figure 1I-1J), when most projection neurons have been generated. As expected, the numbers of red and green wild-type cells were not significantly different at both analysis time-points (Figure 1E and 1J). Next, we generated *Rptor*-MADM [*MADM-11^GT/TG,Rptor^*; *Emx1-CreER^+/-^* (Figure 1F-1H and S1A, see also Methods for details)] clones with red subclones containing wild-type- and green subclones *Rptor^-/-^* mutant cells, respectively. We used the same TM injection/analysis protocol as above and found similar cell numbers like in Control-MADM clones (Figures 1E, 1H, 1J and 1L). Thus, the neurogenic potential of mutant *Rptor^-/-^* and wild-type subclones originating from a symmetrically-dividing RGP was identical.

In the second experiment (Figure 1M-1O), we quantified projection neuron output of individual RGPs upon their switch to asymmetric neurogenic proliferation mode (Gao et al. 2014; Beattie et al. 2020). We injected TM in *Rptor*-MADM at E12 to generate asymmetric MADM clones (Figure 1M-N) and analyzed cell numbers at postnatal day 0 (P0), well after projection neuron generation has been completed. We assessed total clone/unit size, majority population (larger subclone) and minority population (smaller subclone) (Figure 1O). In all three quantifications we could not detect a significant difference in the number of *Rptor^-/-^* mutant cells when compared to *Rptor^+/+^* wild-type. Importantly, the total clone size was about 8-9 neurons for each genotype, which reflects the genuine projection neuron unit of a single neurogenic cortical RGP (Gao et al. 2014; Beattie et al. 2020). The above results might be unexpected, to a predicted reduced neurogenic RGP potential based on the observed microcephaly phenotype in global tissue *Rptor* knockout. Nevertheless, the data from quantitative MADM-based clonal analysis suggests that *Rptor* function is not cell-autonomously required for cortical RGP-mediated projection neuron generation.

### Whole tissue but not sparse ablation of *Rptor* results in microcephaly

The results from MADM-based clonal analysis indicate that at single RGP clone level *Rptor* is not cell-autonomously required for cortical neurogenesis. To corroborate the above analysis and to assess putative non-cell-autonomous effects, we established two additional genetic MADM paradigms in combination with *Emx1*-Cre driver (Gorski et al. 2002) (Figure 2A-2I and S1-S2). First, genetic *Rptor* mosaic (*Rptor*-MADM; *MADM-11^GT/TG,Rptor^*; *Emx1^Cre/+^*) with sparse *Rptor* deletion in just a few RGPs with normal (i.e. heterozygous or wild-type) cortical tissue background. Second, global whole cortex conditional *Rptor* knockout (cKO-*Rptor*-MADM; *MADM-11^GT,Rptor/TG,Rptor^*; *Emx1^Cre/+^*) with *Rptor^-/-^* mutant background (most if not all RGPs mutant for *Rptor*). Thus, individual *Rptor^-/-^* RGPs were either surrounded in a local microenvironment with ‘normal’ cells (Figure 2H, S1A and S2A) or by *Rptor^-/-^* mutant cells (Figure 2I, S1B and S2B). The above MADM paradigms, in comparison with Control-MADM (*MADM-11^GT/TG^*; *Emx1^Cre/+^*) (Figure 2A-B, and 2G), enable the phenotypic analysis at single cell level and therefore provide a highly quantitative platform to assay the influence of the local microenvironment, i.e. non-cell-autonomous effects on *Rptor^-/-^* mutant cells.

**Figure 2.**
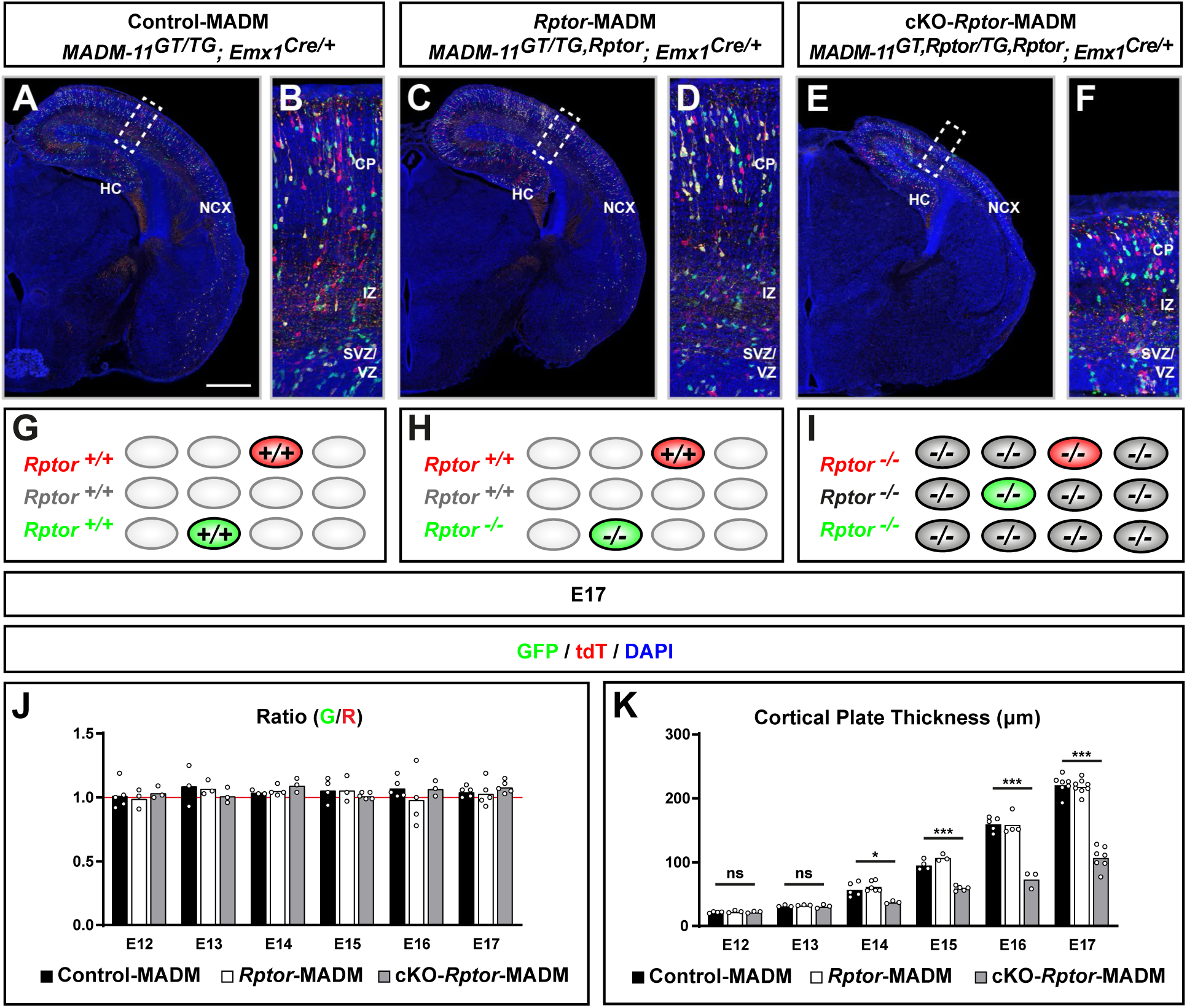
Tissue-wide but not sparse loss of *Rptor* function results in microcephaly. (A-I) Representative images of MADM-labeling pattern in the somatosensory cortex (A-F) and experimental paradigm (G-I) in Control-MADM, GFP^+^ (green), tdT^+^ (red), GFP^+^/tdT^+^ (yellow) and unlabeled (vast majority) cells are all *wild-type* (A, B and G; *MADM-11^GT/TG^*;*Emx1^Cre/+^*); *Rptor*-MADM, GFP^+^ cells are *Rptor^-/-^*, tdT^+^ cells are *Rptor^+/+^*, GFP^+^/tdT^+^ and unlabeled cells are *Rptor^+/-^* (C, D and H; *MADM-11^GT/TG,Rptor^*;*Emx1^Cre/+^*); and cKO*-Rptor*-MADM, all cortical projection neurons (GFP^+^, tdT^+^, GFP^+^/tdT^+^ and unlabeled) are all *Rptor^-/-^* (E, F and I; *MADM-11^GT,Rptor/TG,Rptor^*;*Emx1^Cre/+^*) at E17. Higher magnification (B, D and F) of boxed areas in A, C and E, respectively, illustrating the CP with emerging cortical layers. (J) Quantification of green/red (G/R) ratio of cortical excitatory neurons in the three different MADM paradigms. (K) Embryonic time course analysis (E12-E17) of cortical plate thickness (μm). Nuclei were stained using DAPI (blue). NCX: Neocortex. HC: Hippocampus. Each individual data point represents one experimental animal. Bars represent mean. Significance was determined using two-way ANOVA with Dunnett’s (J) or Tukey’s (K) multiple comparisons. ns: non-significant. *p < 0.05, ***p < 0.001. Scale bar: 500 µm (A, C and E).

We assessed cortical projection neuron generation in time course from E12 (start of cortical neurogenesis) until E17 (when neurogenesis has concluded) in Control-MADM, *Rptor*-MADM and cKO-*Rptor*-MADM (Figure 2J-2K). Since we utilized constitutively active *Emx1*-Cre, MADM labeling events occur stochastically at any given time in a random sparse subset of dividing *Emx1^+^* RGPs, in contrast to the above clonal analysis with defined temporal induction. We found that the ratio of green to red (G/R) cells was approximately one in all three, Control-MADM, *Rptor*-MADM and cKO-*Rptor*-MADM, paradigms and at all sampled time points (Figure 2J). A G/R ratio of ∼1 in Control-MADM and cKO-*Rptor*-MADM was anticipated due to the identical genotypes of red and green cells (both wild-type in Control-MADM and both *Rptor^-/-^* in cKO-*Rptor*-MADM). A ratio of ∼1 in *Rptor*-MADM with wild-type red cells and *Rptor^-/-^* mutant green cells however again indicates that RGP-mediated neurogenesis is not compromised cell-autonomously upon loss of *Rptor*.

Next, we assessed cortical plate thickness at E12-E17 and found that in cKO-*Rptor*-MADM, significant smaller cortical size and thickness was apparent from E14 onward, leading to severe microcephaly at E17 (Figure 2K). Importantly, the microcephaly only emerged in cKO-*Rptor*-MADM (with cortex-wide *Rptor* deletion), but not in mosaic *Rptor*-MADM (with sparse *Rptor* deletion) nor in Control-MADM, suggesting that tissue-wide cell-non-autonomous effects play a dominant role in the emergence of the *Rptor* loss-of-function phenotype in cKO-*Rptor*-MADM.

### Differential gene and protein expression in *Rptor^-/-^* mutant cells upon sparse versus global tissue-wide *Rptor* KO

In order to analyze the consequences of sparse and tissue-wide *Rptor* deletion, and to obtain insight into the nature of cell-autonomous and non-cell-autonomous aberrations at the molecular level we next pursued transcriptome and proteome profiling. We isolated cortical MADM-labeled cells using fluorescent-activated cell sorting (FACS) (Laukoter et al. 2020a; Amberg et al. 2024) from Control-MADM, *Rptor*-MADM and cKO-*Rptor*-MADM at E13, E16 and P0 (Figure 3A). We specifically focused on GFP^+^ cells that were wild-type in Control-MADM, *Rptor^-/-^* in *Rptor*-MADM (normal microenvironment) and cKO-*Rptor*-MADM (*Rptor^-/-^* mutant microenvironment). Next, isolated cells were subjected to RNA sequencing using SMARTer technology and liquid chromatography-tandem mass spectrometry (LC-MS/MS) [see (Hansen et al. 2022) and Methods for details], followed by bioinformatics analysis.

**Figure 3.**
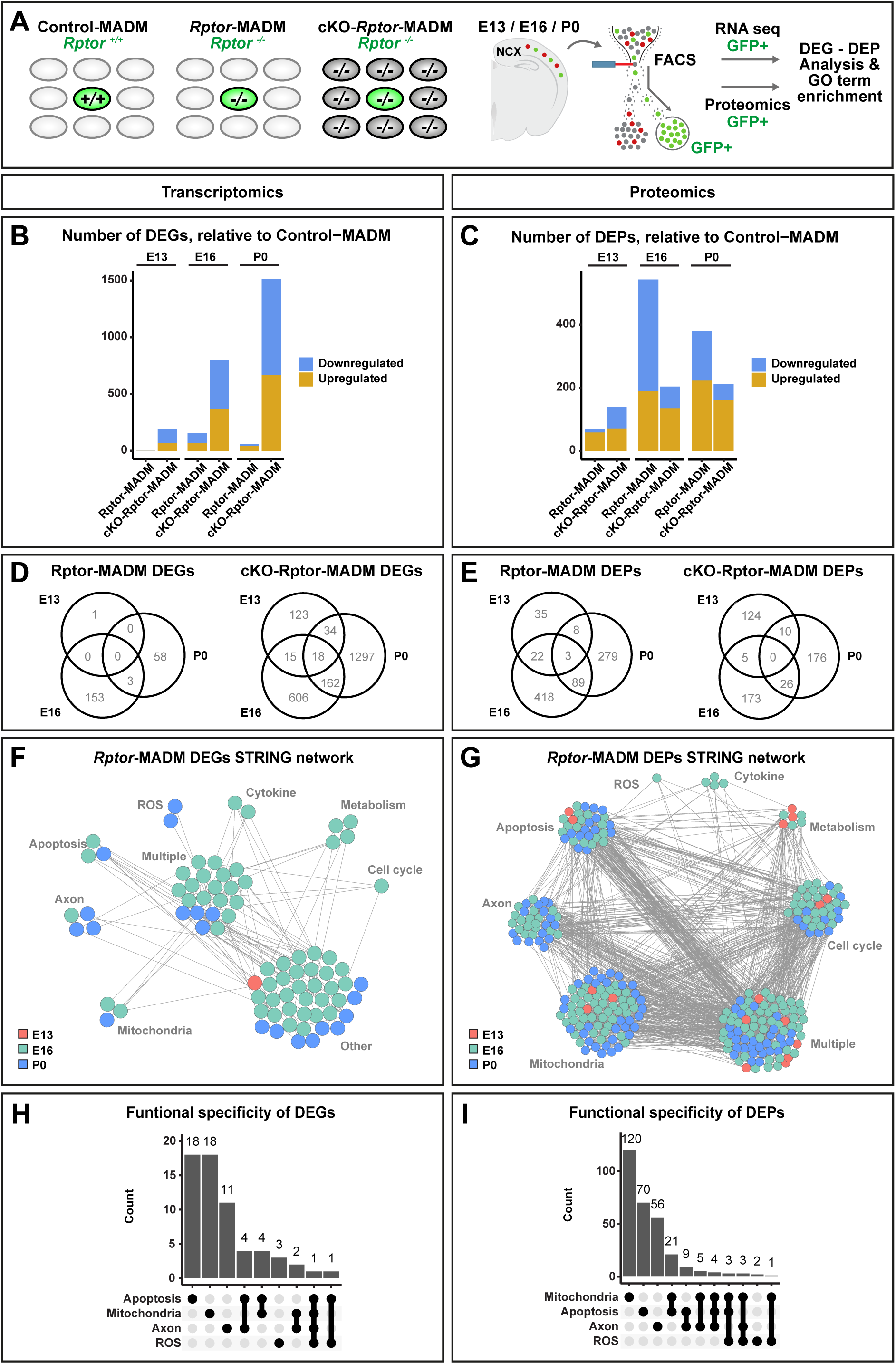
Global tissue-wide and sparse *Rptor* ablation result in distinct transcriptional and proteomic responses. (A) Experimental strategy to FACS-enrich for cortical cells in the *Emx1^+^* lineage. GFP^+^ (green) cells were isolated from Control-MADM (wild-type, *Rptor*^+/+^), *Rptor*-MADM (genetic mosaic, *Rptor*^-/-^) and cKO-*Rptor*-MADM (conditional knockout, *Rptor^-/-^*) at embryonic time points E13, E16 and at P0 for RNA-seq, proteomic assessment, and differential gene/protein expression analysis (DEG/DEP). (B and C) Numbers of differentially expressed genes (B; DEG, FDR < 0.05, DESeq2) and differentially expressed proteins (C; DEPs, FDR < 0.01, t-test) in *Rptor*-MADM and cKO-*Rptor*-MADM relative to Control-MADM at E13, E16 and P0. (D and E) Venn diagrams visualizing the overlap of DEGs (D) and DEPs (E) in *Rptor*-MADM/Control-MADM and cKO-*Rptor*-MADM/Control-MADM across developmental time points. (F and G) Network of 79 DEGs (F) and 285 DEPs (G) determined by STRING. Functional gene groups were predicted by gene ontology. (H and I) Association of DEGs (H) and DEPs (I) with single or different combinations of GO term groups.

First, we focused on the transcriptomic data set and carried out principal component analysis (PCA) across all samples using the top 500 variable genes (Figure S3A). Interestingly, individual samples of all three genetic paradigms clustered closely together at early E13 and E16 timepoints but diverged at P0, indicating an increasing variance in gene expression. Next, we validated *Rptor* deletion in the three paradigms at all sampled time points (Figure S3B). As expected, *Rptor* expression levels were close to zero in *Rptor^-/-^* mutant cells in *Rptor*-MADM and cKO-*Rptor*-MADM when compared to wild-type cells in Control-MADM. Analysis of differentially expressed genes (DEGs) showed massively increased DEG numbers in cKO-*Rptor*-MADM when compared to *Rptor*-MADM (Figure 3B), indicating that tissue-wide *Rptor* ablation leads to much higher deregulation of gene expression than sparse *Rptor* deletion. The vast majority of DEGs displayed temporal (i.e. developmental stage) and genetic ablation paradigm specificity (Figure 3D). Next, we assessed differentially expressed proteins (DEPs) (Figure 3C) in *Rptor*-MADM and cKO-*Rptor*-MADM in comparison to Control-MADM and found that, similar to the DEGs, DEPs exhibit developmental stage and genetic deletion specificity in their deregulated expression profile (Figure 3E). Importantly, DEGs (Figure S3C) and DEPs (Figure S3F) at all sampled timepoints were to a large extent exclusive for the two distinct *Rptor* ablation (global tissue-wide versus sparse) paradigms.

To obtain insight into the biological processes associated with DEGs and DEPs upon *Rptor* deletion we conducted Gene Ontology (GO) term analysis for DEGs (Figure S3D) and DEPs (Figure S3G). We found that a set of cellular processes (metabolism, mitochondria, cell cycle, axon, reactive oxygen, cytokines and apoptosis) that have been previously associated with *Rptor*/*Mtor* were significantly enriched, albeit at different levels across developmental timepoints in both *Rptor*-MADM and cKO-*Rptor*-MADM. Next, we used the Search Tool for the Retrieval of Interacting Genes/Proteins (STRING) to evaluate potential connections or interactions among deregulated cellular processes. Both, DEGs and DEPs formed networks that were significantly more connected than expected at random (DEGs: p=0.0072, DEPs: p<10^-16^). Since DEG and DEP networks contained genes/proteins from different developmental stages and associated with diverse biological functions, these results suggest that molecular phenotypes were connected among themselves and across development (Figure 3F-3I and Figure S3E and S3H). In summary, the sparse and global tissue-wide ablation of *Rptor* lead to significant deregulation of gene/protein expression with common but also highly exclusive sets of DEGs/DEPs with regard to the genetic *Rptor* deletion paradigm and developmental stage.

### Non-cell-autonomous and collective cell-autonomous cellular phenotypes result in overall microcephaly upon tissue-wide loss of *Rptor*

The above gene/protein expression analysis revealed that sparse versus global tissue-wide *Rptor* deletion results in both common and differential gene/protein deregulation in *Rptor^-/-^* mutant cells. Therefore, we next assessed how deregulated gene/protein expression in *Rptor^-/-^* mutant cells translates into actual cellular phenotypes in *Rptor*-MADM and cKO-*Rptor*-MADM that could potentially explain the different brain phenotypes at tissue-wide level (Figure 4).

**Figure 4.**
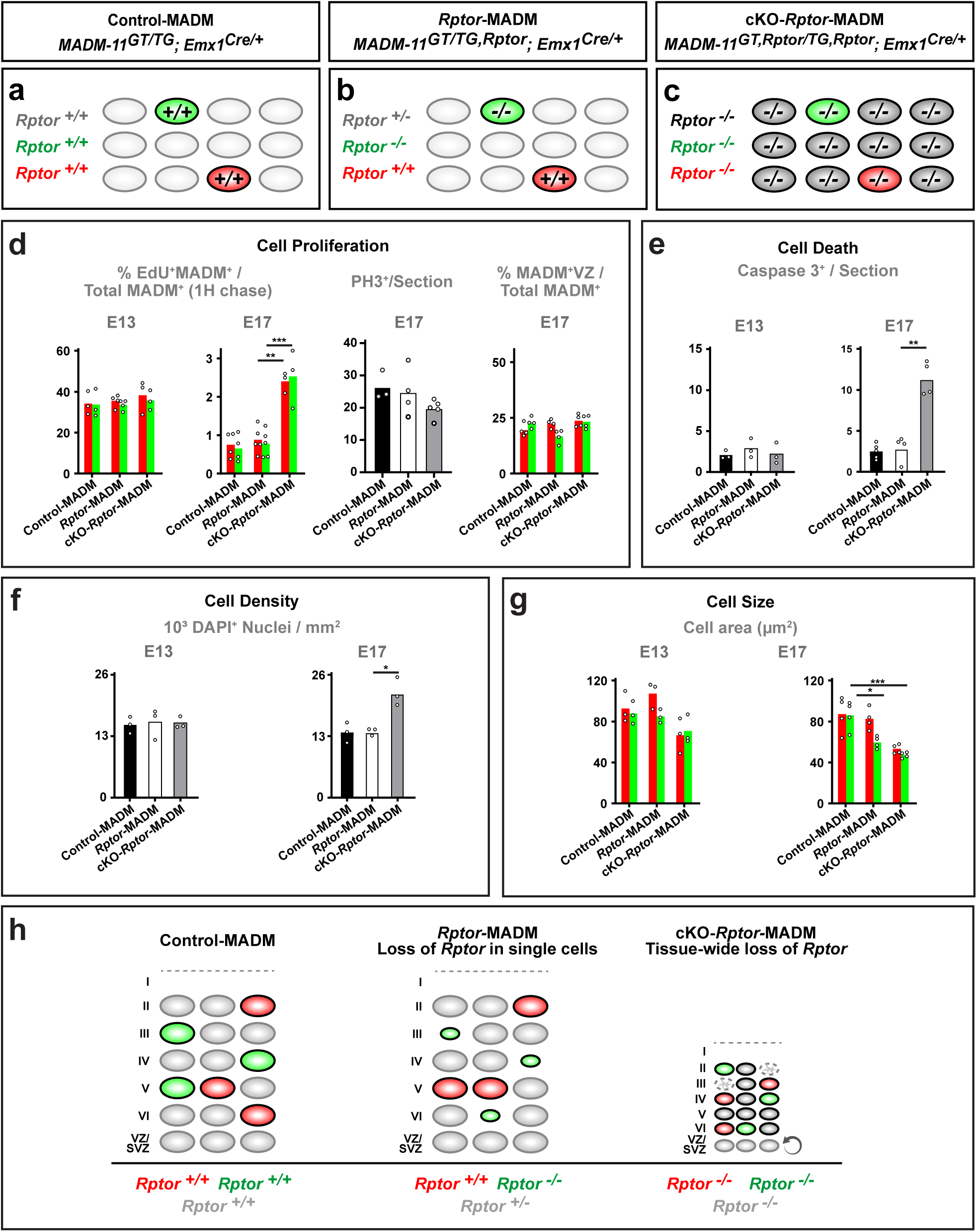
Cumulative non-cell-autonomous phenotypes and collective cell-autonomous decrease in cell size result in overall microcephaly upon tissue-wide loss of *Rptor*. (A-C) Experimental MADM paradigms to assess cell-autonomous *Rptor* function and tissue-wide non-cell-autonomous effects. (A) Control-MADM (*MADM-11^GT/TG^*;*Emx1^Cre/+^*; red cells: *Rptor^+/+^*; green cells: *Rptor^+/+^*; background: *Rptor^+/+^*). (B) *Rptor*-MADM (*MADM-11^GT/TG,Rptor^*;*Emx1^Cre/+^*, red cells: *Rptor^+/+^*; green cells: *Rptor^-/-^*; background: *Rptor^+/-^*). (C) cKO-*Rptor*-MADM (*MADM-11^GT,Rptor/TG,Rptor^*;*Emx1^Cre/+^*), red cells: *Rptor^-/-^*; green cells: *Rptor^-/-^*; background: *Rptor^-/-^*). (D) Quantification of the fraction (%) of EdU^+^/MADM^+^ SVZ/VZ cells of the total MADM-labeled cells at E13 and E17; the fraction (%) of PH3^+^/section and the fraction (%) of MADM^+^ ventricular zone (VZ) cells compared to the total number of MADM^+^ cells at E17. (E) Quantification of the number of Caspase-3^+^ cells per cortical hemisphere at E13 and E17. (F) Quantification of DAPI^+^ nuclei/mm^2^ in the cortical plate. (G) Quantification of the cell area (µm^2^) of MADM^+^ neurons in the cortical plate at E13 and E17. (H) Schematic summarizing the interplay of intrinsic cell-autonomous and tissue-wide contributions in microcephaly formation upon loss of *Rptor*. Each individual data point represents one experimental animal (D, E, F and G). Bars represent mean. Significance was determined using multiple t-test (F and G) or two-way ANOVA with Tukey’s multiple comparisons (D and E). *p < 0.05, **p < 0.01, ***p < 0.001.

First, we analyzed RGP proliferation, since cell-cycle GO terms were significantly enriched in the above gene expression analysis, by injection of EdU 1hour prior fixation at E13 or E17 (Figure 4A-4D). EdU incorporates into newly synthesized DNA during replication, thereby labeling cycling cells in S-phase of the cell-cycle. By fixing the tissue immediately after EdU injection, we obtained a snapshot of cycling cells at respective developmental stages. At early stages of neurogenesis (E13) the percentage of MADM^+^ cells that incorporated EdU was indistinguishable in all three Control-MADM, *Rptor*-MADM and cKO-*Rptor*-MADM. However, by the end of neurogenesis at E17, *Rptor^-/-^*mutant cells in cKO-*Rptor*-MADM showed increased incorporation of EdU, while wild-type cells in Control-MADM and both *Rptor^-/-^* (green bars in Figure 4D) and *Rptor^+/+^* (red bars in Figure 4D) cells in mosaic *Rptor*-MADM showed similar - overall lower - EdU incorporation rates (Figure 4D). Next, we stained against the M-phase marker PH3 but did not observe a significant difference in the number of PH3^+^ cells across all three MADM paradigms (Figure 4D). We also quantified the number of MADM^+^ cells in the VZ, as a proxy for the number of progenitor cells, but did not observe any difference between the three MADM paradigms (Figure 4D). Together, the above results suggest that *Rptor* ablation leads to a modification of RGP cell cycle length at late stages of neurogenesis without apparent changes in proliferation rate *but* only in whole tissue *Rptor* deletion context and not upon sparse *Rptor* deletion, demonstrating non-cell-autonomous phenotypic origin. Second, we quantified cell death in cortical tissue at E13 and E17 upon sparse and global tissue-wide *Rptor* ablation in *Rptor*-MADM and cKO-*Rptor*-MADM paradigms, respectively. We performed immunostainings against activated Caspase-3 as proxy for apoptotic cells. We did not observe a significant difference in the number of Caspase-3^+^ cells per section between Control-MADM and *Rptor*-MADM or cKO-*Rptor*-MADM, respectively, at E13 (Figure 4E). In contrast, the number of Caspase-3^+^ cells were significantly increased at E17 upon global tissue-wide *Rptor* deletion in cKO-*Rptor*-MADM but not in *Rptor*-MADM with sparse *Rptor* deletion when compared to Control-MADM (Figure 4E).

Third, we assessed overall cell density (i.e. DAPI^+^ Nuclei / mm^2^). While cell density was similar in Control-MADM, *Rptor*-MADM and cKO-*Rptor*-MADM at E13, we detected a significant increase in cell density in cKO-*Rptor*-MADM at E17 but not in *Rptor*-MADM when compared to Control-MADM (Figure 4F).

Lastly, we measured cell size – which is well known to be regulated by RPTOR/mTORC1 signaling (Liu and Sabatini 2020; Battaglioni et al. 2022). At E13, cell size was comparable in *Rptor^+/+^*and *Rptor^-/-^*cells across Control-MADM, *Rptor*-MADM and cKO-*Rptor*-MADM paradigms (Figure 4G). However, at E17 we observed a significant decrease in cell size in *Rptor*^-/-^ cells when compared to *Rptor^+/+^* cells in mosaic *Rptor*-MADM (Figure 4G). Thus, the sparse elimination of *Rptor* resulted in diminished cell size which reflects a cell-autonomous loss-of-function phenotype. In cKO-*Rptor*-MADM we detected decreased cell size in *Rptor^-/-^*cells that was similar in scale to the decrease in *Rptor^-/-^* cells in mosaic *Rptor*-MADM (Figure 4G).

Based on the above results we conclude that 1) cell-non-autonomous effects emerging as a result of global tissue-wide *Rptor* ablation in cKO-*Rptor*-MADM led to increased cell-cycle length in late neurogenic RGPs, higher amounts of apoptotic cells and increased cell density in the developing *Rptor^-/-^*mutant cortex; 2) decreased cell size reflects a true cell-autonomous phenotype upon both sparse and global tissue-wide *Rptor* deletion. Altogether, our data suggest that the observed microcephaly upon global tissue-wide *Rptor* elimination emerges as a result of adverse cumulative non-cell-autonomous effects in combination with collective cell-autonomous decrease in cell size, rather than due to cell-autonomous deficits in RGP lineage progression and cortical projection neuron generation (Figure 4H).

### Cell-autonomous *Rptor* function is required for postnatal cortical projection neuron maintenance

Global brain/tissue-wide *Rptor* deletion results not only in microcephaly but also leads to death at, or within a few days after birth (Cloetta et al. 2013). Whether *Rptor* fulfills a role in postnatal cerebral cortex development is therefore currently unknown. The sparse mosaic *Rptor* deletion paradigm, with only a very small fraction of *Rptor^-/-^* mutant projection neurons, can bypass early postnatal lethality and thus enabled the more long-term functional analysis of *Rptor* in cortical development. In a first pass analysis we assessed cortical projection neuron numbers in a time course. We analyzed the ratio of green/red cells from P0 up to 12 months (Mo) in Control-MADM, both green and red cells with *Rptor^+/+^* genotype; and *Rptor*-MADM, red cells with *Rptor^+/+^* and green cells with *Rptor^-/-^* genotype (Figure 5A-5K). As expected, the G/R in Control-MADM was ∼1 at all analyzed timepoints. In contrast, the G/R ratio was significantly decreased in *Rptor*-MADM from birth onward, with less than 50% of *Rptor^-/-^*mutant cells after the first postnatal week (Figure 5I). The G/R ratio was decreased across all cortical layers with a slightly stronger decrease in layers II-IV in *Rptor*-MADM (Figure 5I-5K). We also observed decreased numbers of *Rptor^-/-^* mutant projection neurons with either maternal or paternal disomy (Laukoter et al. 2020a; Pauler et al. 2021) (Figure S4A-S4H), and when mutant *Rptor^-/-^* cells were labeled with tdT (red) instead of GFP (green) (Figure S4I-S4P).

**Figure 5.**
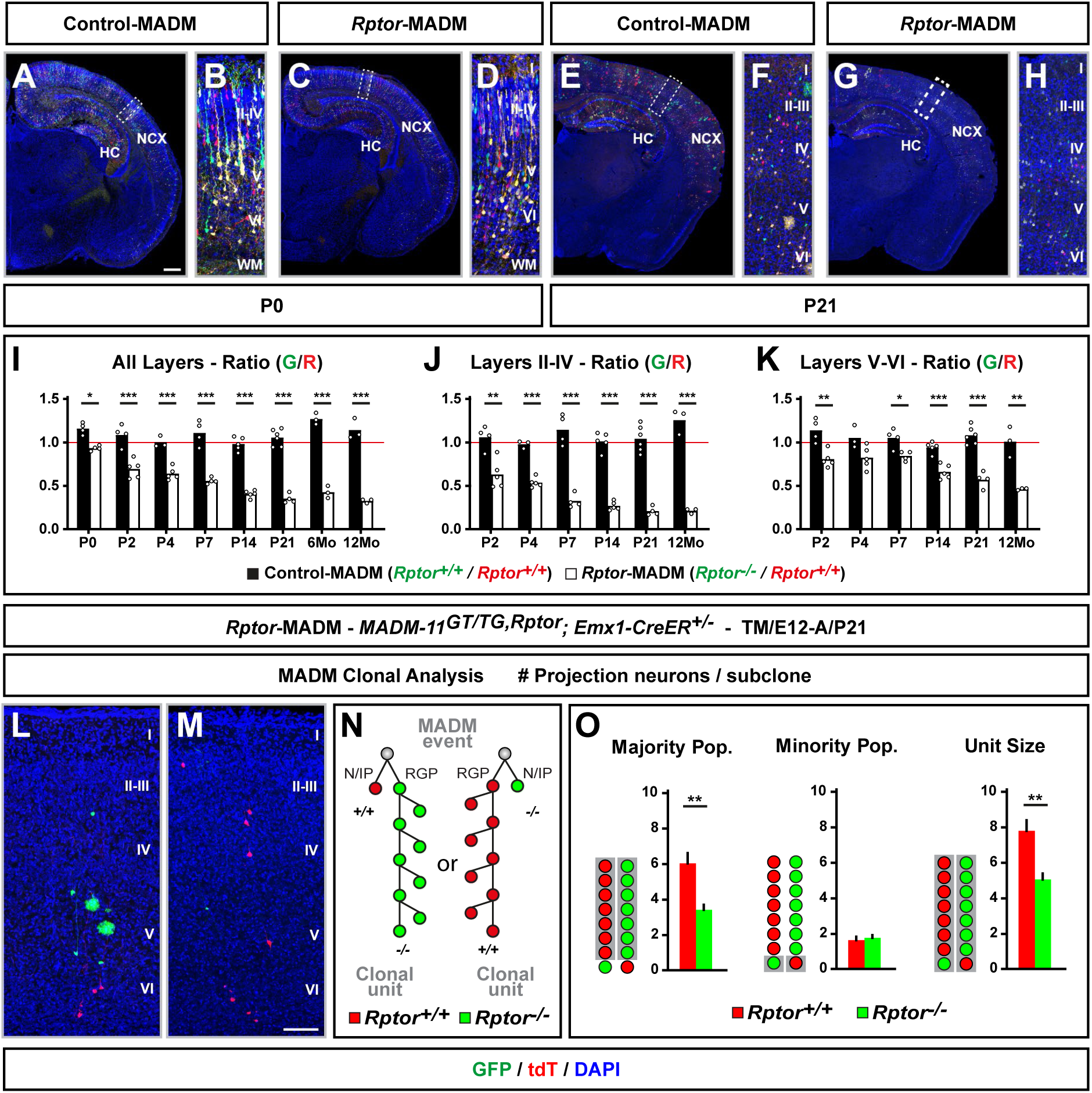
*Rptor* is cell-autonomously required for postnatal projection neuron maintenance. (A-H) Representative images of MADM-labeled cortical projection neurons in Control-MADM (A, B, E and F, *MADM-11^GT/TG^*;*Emx1^Cre/+^*; red cells: *Rptor^+/+^*; green cells: *Rptor^+/+^*; background: *Rptor^+/+^*); and *Rptor*-MADM (C, D, G and H, *MADM-11^GT/TG,Rptor^*;*Emx1^Cre/+^*; red cells: *Rptor^+/+^*; green cells: *Rptor^-/-^*; background: *Rptor^+/-^*) at P0 (A-D) and P21 (E-H). (I-K) Time course analysis of G/R ratio of MADM-labeled cortical projection neurons in all cortical layers (I), layers II-IV (J), and layers V-VI (K) at P0, P2, P4, P7, P14, P21, 6 months (6Mo), and 1 year (12Mo). (L-O) MADM clonal analysis in *Rptor*-MADM (*MADM-11^GT/TG,Rptor^*;*Emx1-CreER^+/-^*) with TM injection at E12 and analysis at P21 to determine the maintenance of the clonal projection neuron unit in asymmetric neurogenic clones. Note that the clonal unit output was unchanged up to P0 as indicated in Figure 1. (L and M) 3D reconstruction images of representative asymmetric G2-X MADM clones with majority population in green (L) or red (M) in *MADM-11^GT/TG,Rptor^ Emx1-CreER^+/-^*. (N) Schematic illustration of the MADM clonal analysis paradigm. (O) Quantification of the size of the majority population, minority population and the unitary size of asymmetric neurogenic clones in *Rptor*-MADM samples at P21. Nuclei were stained using DAPI (blue). NCX: Neocortex. HC: Hippocampus. Each individual data point represents one experimental animal (I, J and K). Bars and error bars represent mean ± SEM. Significance was determined using unpaired *t* test (O) or multiple unpaired *t* test (I and J) or two-way ANOVA with Šídák’s multiple comparisons (K). *p < 0.05, **p < 0.01, ***p < 0.001. Cortical layers are indicated in roman numerals. Scale bar: 500 µm (A, C, E, and G), 50 µm (L and M).

Next, we assessed individual mosaic MADM clones, in *MADM-11^GT/TG,Rptor^*; *Emx1-CreER^+/-^* with red *Rptor^+/+^* and green *Rptor^-/-^* cells, induced at the start projection neuron generation at E12 and analyzed at P21 (Figure 5L-5O). While the genuine projection neuron unit size in MADM-labeled cortical clones measures about 8-9 neurons in total (Gao et al. 2014; Llorca et al. 2019), the total unit size was significantly reduced in mosaic *Rptor*-MADM clones (Figure 4O). Thus, while mosaic *Rptor*-MADM clones were found to contain 8-9 neurons at P0 (see Figure 1O), the unit size decreased significantly during the first postnatal weeks. Together, the above data from sparse *Rptor* deletion at population (Figure 5A-5K) and individual clone (Figure 5L-5O) level indicate that *Rptor* is cell-autonomously required for postnatal projection neuron maintenance.

To corroborate the above finding we next assessed the numbers MADM-labeled cortical projection neurons in *Mtor*-MADM, using MADM-4 reporters (Contreras et al. 2021) in combination with *Emx1*-Cre (Gorski et al. 2002), with red *Mtor^+/+^* and green *Mtor^-/-^* cells at E16, P0 and P16 (Figure S5). Like in *Rptor*-MADM, projection neuron generation was unaffected with G/R of ∼1 at E16 in *Mtor*-MADM. Yet, postnatal maintenance was similarly affected in the absence of *Mtor* mirroring the findings upon *Rptor* deletion Figure S5M).

### Single-cell RNA sequencing reveals cell-type-specific *Rptor* function across distinct cortical projection neuron classes

*Rptor* function appears to be essential for postnatal projection neuron maintenance, and layer II-IV projection neurons showed somewhat increased propensity to be lost when compared to neurons in layers V-VI. In order to assess putative cell-type specificity of *Rptor* function at the molecular level we next pursued single-cell RNA sequencing (scRNA-seq) (Figure 6A). We chose two developmental stages, E18 and P1, marking the onset of neuron loss upon *Rptor* deletion, and isolated green GFP^+^ MADM-labeled cells from Control-MADM (*Rptor^+/+^*) and *Rptor*-MADM (*Rptor^-/-^*) by using FACS. Samples were processed using 10X Genomics Technology and subjected to scRNA-seq (see Methods for details). After quality control, we recovered 10612 and 9365 cells, at E18 and P1, respectively, for downstream bioinformatics analysis. Cells that could unambiguously assigned to either *Rptor^+/+^* or *Rptor^-/-^*genotype were of similar quality (Figure S6A-S6B). Uniform manifold approximation and projection (UMAP) in combination with unsupervised clustering, label transfer of cell types from a reference data set (Di Bella et al. 2021), and marker gene expression could identify all major cell types of the *Emx1^+^* cell lineage (Figure S6C-S6H). Next, we focused the analysis on postmitotic excitatory projection neurons (5269/2148 cells at E18/P1; Figure 6B) and classified cells into *Satb2*-expressing callosal projection neurons (CPNs) and *Ctip2*-expressing subcerebral projection neurons (SCPNs) (Figure 6C and S6I). The relative abundance of *Satb2^+^*/*Ctip2^+^* cells did not change upon loss of *Rptor* (Figure S6J).

**Figure 6.**
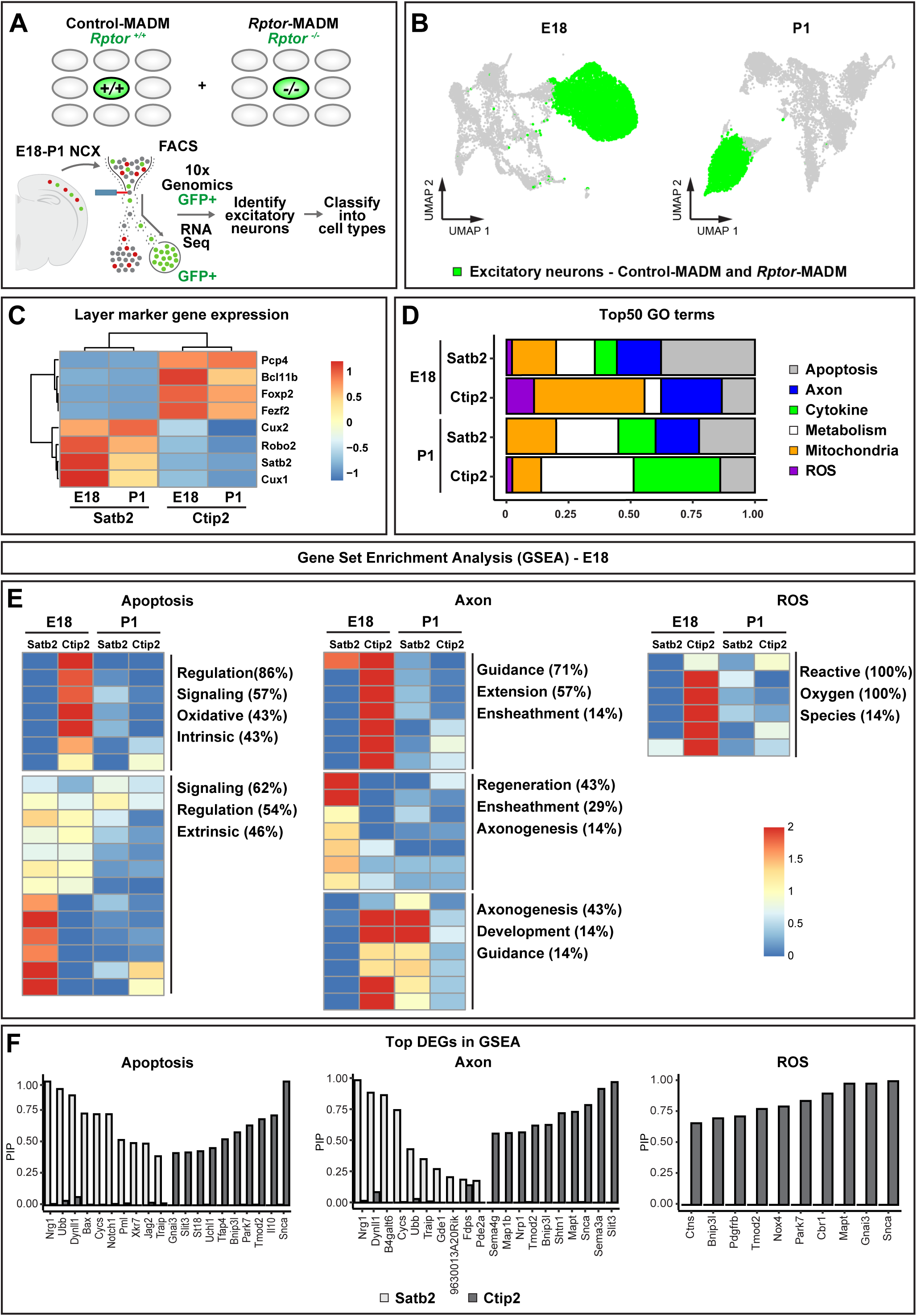
Single cell RNA sequencing reveals differential cell-type specific requirements for *Rptor* function across distinct projection neuron classes. (A) Strategy to FACS-enrich cortical cells in the *Emx1*^+^ lineage, and to identify different classes of excitatory projection neurons. GFP^+^ (green) cells were isolated from Control-MADM (*Rptor^+/+^*) and *Rptor*-MADM (*Rptor^-/-^*) at E18 and P1, processed for 10x Genomics, RNA sequencing and bioinformatics analysis. (B) UMAPs displaying 10612 and 9365 cell pools from E18 and P1, respectively. Excitatory neurons were labeled in green. (C) Pseudo-bulk expression of marker genes in *Ctip2*- and *Satb2*-expressing neuron populations. (D) *Rptor*-MADM versus Control-MADM DEGs at E18 and P1 in *Ctip2*- and *Satb2*-expressing neuron populations were subjected to integrative differential expression and gene set enrichment analysis (iDEA) using selected gene ontology (GO) terms. Bars indicate fractions of GO groups within top 50 GO terms identified in each sample. (E) Heatmaps of GO term p-value scores in *Ctip2*- and *Satb2*-expressing neuron populations at E18 and P1. To enhance readability, fractions of top terms in GO term descriptions were depicted. (F) Posterior inclusion probabilities (PIP) of top DEGs in GO terms in *Ctip2*- and *Satb2*-expressing neuron populations at E18.

To gain insight into the transcriptional changes and associated biological processes due to loss of *Rptor* we next pursued integrative differential expression and gene set enrichment analysis [iDEA; (Ma et al. 2020)]. In a first pass analysis, we extracted the top 50 enriched GO terms, grouped them, and plotted their relative abundance (Figure 6D). Specific cell biological processes (apoptosis, axon, cytokine, metabolism, mitochondria, reactive oxygen species), previously associated with RPTOR/mTORC1 function were significantly deregulated in both *Satb2^+^*CPNs and *Ctip2^+^* SCPNs at E18 and P1, albeit at varying levels (Figure 6D). Next, we assessed the cell-type specificity of actual GO terms and found that defined GO terms showed differences in enrichment not only between *Satb2^+^* CPN and *Ctip2^+^* SCPN cell-types but also across developmental stages (Figure 6E and S7). Plotting the posterior inclusion probability of genes associated with respective GO term groups that were identified above, revealed highly cell-type-specific gene expression changes upon loss of *Rptor* at E18 (Figure 6F). Altogether, the analysis of deregulated gene expression in *Rptor^-/-^* mutant cells indicated that certain cellular processes were affected in both *Satb2^+^* CPNs and *Ctip2^+^*SCPNs, but that some processes might be affected in a more cell-type specific manner.

To assess how predicted changes in gene expression translate into cellular phenotypes we first focused on axonal projection patterns. To this end we calculated the volume ratio of green GFP^+^ projections - *Rptor^+/+^*in Control-MADM and *Rptor^-/-^* in *Rptor*-MADM - to red *Rptor^+/+^* control projections after eliminating yellow (*Rptor^+/-^*) cells and projections from the analysis (see Methods for details). Cortico-thalamic projection volume ratio was significantly reduced by two-thirds (Figure S8A-K) while the callosal projection volume ratio was reduced by more than half in *Rptor*-MADM when compared to Control-MADM (Figure S8L-T).

While apoptosis and axon-related GO terms were found in both *Satb2^+^* CPN and *Ctip2^+^* SCPN cell-types, albeit with variable enrichment, ROS (reactive oxygen species)-related GO terms were exclusively found in *Ctip2^+^* SCPN cell-types (Figure 6F, right). Therefore, we next assessed ROS production *in situ* at the cellular level. We isolated cortical tissue from Control-MADM and *Rptor-MADM* at P0 (before cell loss) and P2 (shortly after significant cell death), disaggregated the cells, stained with CellROX^TM^ (fluorogenic probe to measure oxidative stress in live cells), and isolated green GFP^+^ single cells by FACS (Figure S9A). We then quantified GFP^+^/ROS^+^ over total GFP^+^ cells and measured mean CellROX intensity. While at P0 we could not observe a significant difference between Control- and *Rptor*-MADM, both parameters were significantly increased at P2 (Figure S9B-S9C). Next, we isolated GFP^+^/ROS^+^ and GFP^+^/ROS^-^ cells to pursue bulk RNA-seq and determine the relative abundance of *Satb2^+^* CPN and *Ctip2^+^* SCPN cell-types within the respective samples. While *Satb2^+^* CPN were equally present, *Ctip2^+^* SCPN cell-types were significantly enriched in the GFP^+^/ROS^+^ but depleted in GFP^+^/ROS^-^ cell populations (Figure S9D).

Altogether, the above scRNA-seq analysis and observations *in situ* provide evidence that *Rptor*/mTORC1 signaling exerts certain critical cellular functions in both *Satb2^+^*CPN and *Ctip2^+^* SCPN cell-types, but also that *Rptor* function shows high level of cell-type specificity.

### *Rptor* promotes cell-type-specific cortical projection neuron survival in *Bax*-dependent and *Bax*-independent manner

Cortical projection neurons across all layers, including *Satb2^+^* CPN and *Ctip2^+^* SCPN cell-types, show high degree of vulnerability and a large fraction thereof is lost, presumably due to apoptosis, upon loss of *Rptor* function. To obtain mechanistic insight at the molecular level we next conceived genetic interaction, i.e. rescue, experiments in MADM context. We first focused on the p53 signaling pathway since deletion of *Trp53* has been shown to abolish apoptosis of cortical projection neurons, in certain contexts (Amberg et al. 2022). Conveniently, the *Trp53* gene is located on the same MADM reporter chromosome (chr.11) like *Rptor*. We thus could generate double mutant *Rptor^-/-^*;*Trp53^-/-^* cells in mosaic context with ‘normal’ background (i.e. double heterozygote or wild-type) (see Methods for details and Figure S10A). However, when we evaluated the G/R ratio in *Rptor*-*Trp53*-MADM (*MADM-11^GT/TG,Rptor,Trp53^*;*Emx1^Cre/+^*) where green GFP^+^ cells were *Rptor^-/-^*;*Trp53^-/-^* and red tdT^+^ cells *Rptor^+/+^*;*Trp53^+/+^* cells, we observed a similar reduction of mutant cells like in *Rptor*-MADM (Figure 7A-7I and 7M). Thus, concomitant deletion of *Trp53* and *Rptor* cannot rescue cortical projection neurons from apoptosis (Figure 7N).

**Figure 7.**
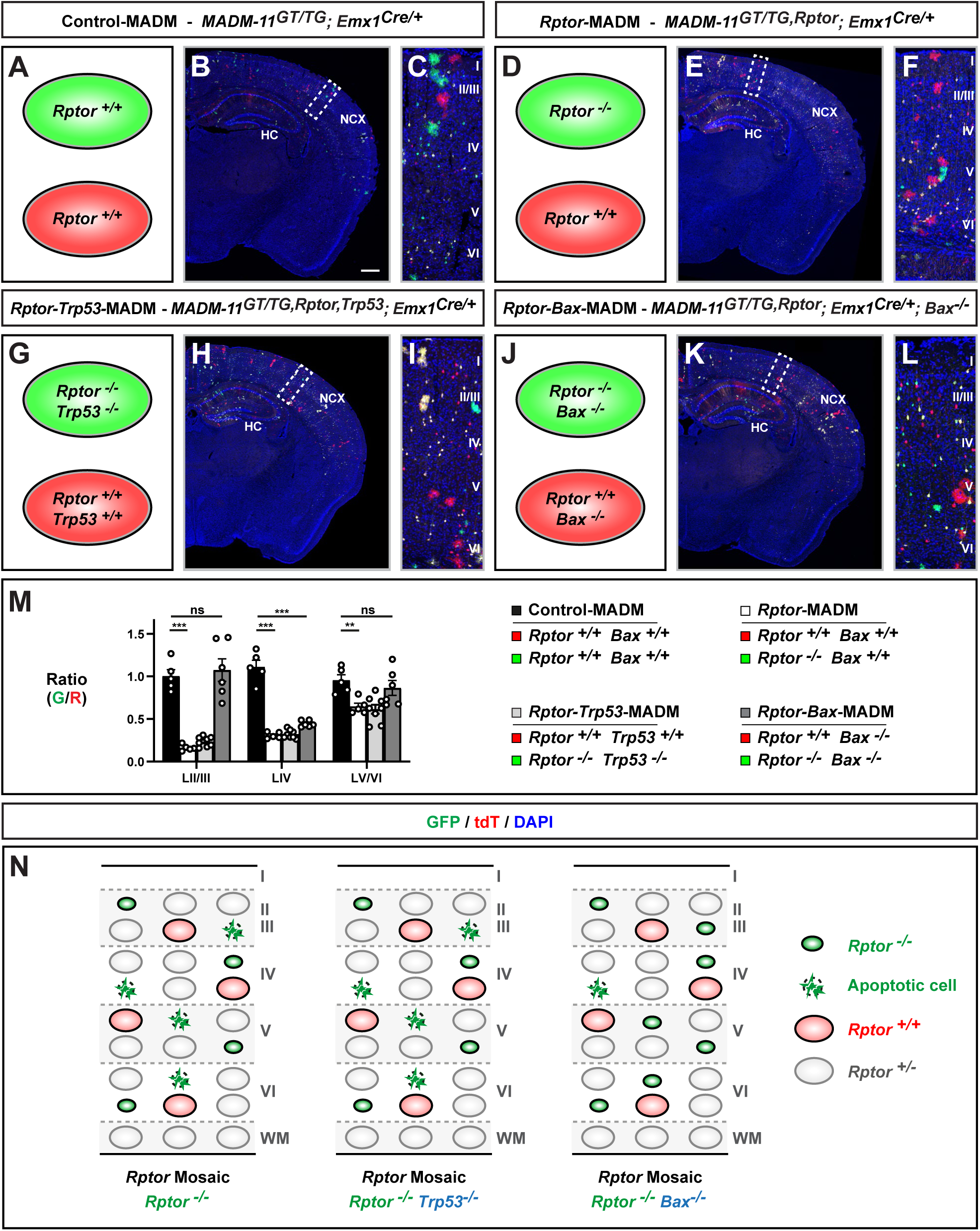
*Rptor* function is required for postnatal projection neuron survival in a cell-type-specific manner. (A-M) Analysis of MADM-labeled projection neurons in Control-MADM (A-C, *MADM-11^GT/TG^*;*Emx1^Cre/+^*; all cells *wild-type*); *Rptor*-MADM (D-F, *MADM-11^GT/TG,Rptor^*;*Emx1^Cre/+^*; red cells: *Rptor^+/+^*; green cells: *Rptor^-/-^*; background: *Rptor^+/-^*); *Rptor-Trp53*-MADM (G-I, *MADM-11^GT/TG,Rptor,Trp53^*;*Emx1^Cre/+^*; red cells: *Rptor^+/+^*;*Trp53^+/+^*; green cells: *Rptor^-/-^*;*Trp53^-/-^*; background: *Rptor^+/-^*;*Trp53^-/-^*); *Rptor-Bax*-MADM (J-L, *MADM-11^GT/TG,Rptor^*;*Emx1^Cre/+^*;*Bax^-/-^*; red cells: *Rptor^+/+^*;*Bax^-/-^*; green cells: *Rptor^-/-^*;*Bax^-/-^*; background: *Rptor^+/-^*;*Bax^-/-^*) with the schematic representation of the respective genotype indicated (A, D, G and J). (B, E, H, and K) Overview of MADM-labeling pattern in the somatosensory cortex of each MADM paradigm at P21. (C, F, I and L) Higher magnification of boxed areas in B, E, H and K, respectively, illustrating the distinct cortical layers. (M) Quantification of G/R ratio of projection neurons in each cortical layer for each genotype. Note that in *Rptor-Bax*-MADM, a rescue is observed in the number of projection neurons for layers II/III and V/VI but not layer IV. No rescue in *Rptor-Trp53*-MADM indicates that cell death occurs in a *Trp53* independent manner. (N) Schematic summarizing *Rptor*/*Trp53* and *Rptor*/*Bax* epistasis experiments and illustrating *Trp53*-independent but cell type specific *Bax*-dependent postnatal cell death upon loss of *Rptor*. Nuclei were stained using DAPI (blue). NCX: Neocortex. HC: Hippocampus. Each individual data point represents one experimental animal. Bars and error bars represent mean ± SEM. Significance was determined using two-way ANOVA with Tukey’s multiple comparisons (M). ns: non-significant. **p < 0.01, ***p < 0.001. Cortical layers are indicated in roman numerals. Scale bar: 500 µm (B, E, H and K).

Next, we focused on the pro-apoptotic gene *Bax* that is well known to trigger apoptosis. The *Bax* gene is located on chr.7 and we therefore generated *Bax* cKO in combination with *Rptor*-MADM (*MADM-11^GT/TG,Rptor^*;*Emx1^Cre/+^*;*Bax^flox/flox^*) (Figure 7J-7L and S10B). In such genetic context – sparse deletion of *Rptor* in *Bax^-/-^*background – we observed that the G/R ratio was ∼1 and not significantly different from G/R ratio in Control-MADM, in cortical layers II/III and V/VI but not in layer IV (Figure 7M). Thus, co-deletion of *Bax* in *Rptor^-/-^* mutant cells could rescue certain cortical projection neuron populations in cell-type-specific manner (Figure 7N).

## DISCUSSION

The generation of a correctly-sized cerebral cortex during development is a highly dynamic and complex morphometric process. NSCs generate large numbers of new neurons every second for a defined period of time, which can be days such as in rodents or months like in human. The proliferation behavior and NSC lineage progression must be tightly regulated in order to generate the appropriate amount and level of neuron diversity that in the end form the mature cerebral cortex. The genetic cues regulating NSC behavior are diverse in nature and depending on the cellular function, compromising single-point mutations in specific genes can lead to cortical malformation such as microcephaly (Pirozzi et al. 2018). Here we have assessed the consequences upon sparse single cell or tissue-wide genetic ablation of *Rptor*, encoding a component of the mTORC1 complex. While global tissue-wide *Rptor* deletion results in severe microcephaly, *Rptor* function is not required cell-autonomously for RGP-mediated cortical projection neuron generation. Rather, a combination of 1) collective decrease in projection neuron cell size and 2) non-cell-autonomous tissue-wide effects underlie the etiology of microcephaly in *Rptor* conditional cortex-specific knockout (Figure 8A). Intriguingly, *Rptor* function is critical at later postnatal stages to safeguard projection neurons from undergoing apoptosis, in a cell-type specific manner (Figure 8B). Below we focus our discussion on the role of *Rptor*/mTORC1 in 1) controlling neurogenesis and cortex size; and 2) generating cortical cell-type diversity and cell-type specificity of *Rptor*/mTORC1 function in cortical development.

**Figure 8.**
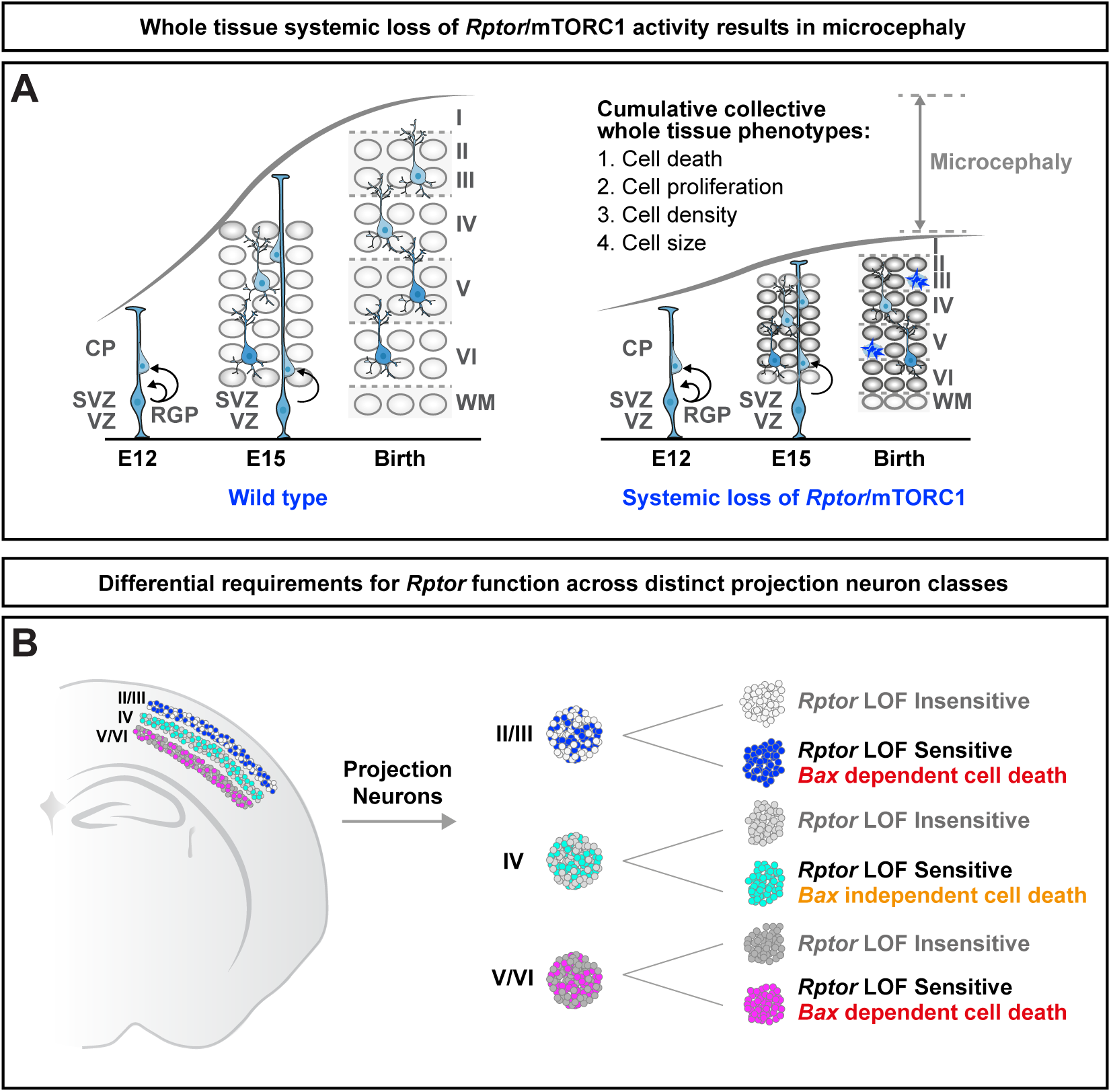
*Rptor* function prevents cortical microcephaly at the global tissue-level and cell-autonomously protects cortical projection neurons in a cell-type specific manner. (A) Schematic model of cortical development in wild-type and upon global whole tissue removal of *Rptor*/mTORC1 activity. The observed microcephaly phenotype upon conditional tissue-wide loss of *Rptor* is due to cumulative collective whole tissue phenotypes rather than due to a deficit in cortical neurogenesis. (B) *Rptor* function is cell-autonomously required for postnatal projection neuron survival in a highly cell-type specific manner. In each cortical layer there are neurons that survive regardless of *Rptor* presence or loss-of-function (LOF), and populations of projection neurons that are sensitive (i.e. undergo apoptosis) to the loss of *Rptor* function. The sensitivity to *Rptor* LOF is not only cell-type specific but projection neurons in layers II/III and V/VI but not IV undergo apoptosis in *Bax*-dependent manner.

### Role of *Rptor*/mTORC1 in controlling neurogenesis and cerebral cortex size

The control of appropriate brain size and prevention of microcephaly during development requires tight regulation of RGPs at the level of gene expression and proliferation behavior. Although more than thirty genes have to date been identified and catalogued in OMIM (Amberger et al. 2019), that when mutated in human cause autosomal recessive primary microcephaly (MCPH), components of the mTORC1 pathway cannot be found in the MCPH gene list. However, the loss of mTORC1 activity by conditional knockout in mice has been shown to result in severe microcephaly (Cloetta et al. 2013; Ka et al. 2014) at the macroscopic level. Yet, how the loss of mTORC1 activity translates to deficits in cortical projection neuron generation with single cell resolution remains unclear. Through MADM-based quantitative lineage tracing, we have previously established a quantitative framework of RGP lineage progression with predictable cortical projection neuron output at any given time throughout embryonic development (Lin et al. 2021; Llorca and Marin 2021; Hippenmeyer 2023). Hence, eliminating *Rptor*/mTORC1 in individual RGPs, and their clonal units of cortical projection neurons, allowed precise determination at which stage and to what extent mTORC1 is required for cortical neurogenesis. Perhaps surprisingly however, by using clonal analysis, we could demonstrate that *Rptor* does not cell-autonomously regulate cortical projection neuron generation (and thus overall neuron number) at the level of individual RGPs although global tissue-wide ablation of *Rptor* leads to dramatic microcephaly - an apparent conundrum? Based on the above findings non-cell-autonomous mechanisms and/or tissue-wide effects clearly seem critical in the emergence of *Rptor* mutant cortex size phenotype. Indeed, when sparse single cell assays were contrasted with global tissue-wide removal of *Rptor*, the interplay of cell-autonomous *Rptor*/mTORC1 function in combination with non-cell-autonomous effects became evident. Based on RNA sequencing and cellular assays, the global tissue-wide, but not sparse/clonal, elimination of *Rptor* let to alteration in RGP cell cycle and increased apoptosis, in agreement with earlier reports (Cloetta et al. 2013; Ka et al. 2014). Interestingly, tissue-wide elimination of *Rictor*, which encodes an essential component of mTORC2 complex, did not result in microcephaly or thinner neocortex despite slightly decreased cell size of neurons (Carson et al. 2013). Thus, the control of macroscopic brain size appears to be controlled mainly by mTORC1.

In conclusion, a collective decrease in cell size, which reflects a true cell-autonomous mTORC1 function, and cumulative non-cell-autonomous effects results in the macroscopic appearance of cortical microcephaly upon loss of mTORC1 function. It will be important in future studies to systematically dissect the interplay of cell-autonomous gene function and global tissue-wide effects in determining brain size not only in mice but also with respect to putative human/species–specific aspects. Perhaps interestingly, although human ‘mTORopathies’ such as hemimegalencephaly or FCD emerge from aberrant mTOR over-activation (Crino 2011; Magri and Galli 2013; Pirozzi et al. 2018; Blumcke et al. 2021; Bizzotto and Walsh 2022; Girodengo et al. 2022), there are currently no reports on human microcephaly patients with loss of function mutations in *RPTOR*/*MTORC1*. However, hypoactivation of the mTOR pathway was recently shown to constitute a converging molecular mechanism that contributes to the clinical symptoms of genetically distinct lissencephaly disorders (Zhang et al. 2025). Thus, mTORC1 activity appears to not only control brain size but also other critical aspects of cerebral cortex development.

### Generating cortical cell-type diversity and cell-type specificity of *Rptor*/mTORC1 function in cortical development

The mechanisms generating cortical projection neuron diversity are not well understood (Greig et al. 2013; Oberst et al. 2019; Di Bella et al. 2024; Pipicelli et al. 2025). While RGPs generate distinct classes of projection neurons in a progressive temporal order (i.e. lower layer neurons first, followed by upper layers), the molecular mechanisms, and in particular the role of mTORC1 signaling in driving RGP lineage progression and temporal neurogenic fate potential are not clear. Interestingly, recent studies that focused on members of the TSC (tuberous sclerosis complex) negative regulator of mTOR signaling, have shown that developmental deletion of TSC proteins (with concomitant increase of mTORC1 activity) at global tissue scale altered RGP and IP balance (Casingal et al. 2025). These deficits translated to changes of clonal unit composition with increased upper layer neuron generation and cortical connectivity. However, we have shown here that endogenous *Rptor*/mTORC1 is not per se required for cortical neurogenesis. In contrast, the maintenance of distinct classes of projection neurons critically relies on mTORC1. Upon *Rptor*/*Mtor* loss of function a defined fraction of projection neurons across all cortical layers was lost due to cell death. Yet, not all layers were equally affected but more neurons in upper layers were lost than in lower layers, indicating cell-type specificity of the sensitivity to loss of mTORC1 function. The mechanisms associated with projection neuron cell death were *Trp53*-independent but included *Bax*-dependent and independent apoptosis pathways. Interestingly, both callosal- and subcortical projection neurons showed axonal projection phenotypes, proceeding cell loss, which could reflect a consequence of the programmed cell death or the cause of it. Although previous studies have associated mTORC1 with axon growth, guidance and regeneration (Park et al. 2008; Terenzio et al. 2018), the evaluation of the precise cell-type and developmental stage-specific requirements and features of mTORC1 signaling await future studies.

A first step towards deeper insight constitutes our single cell gene expression data set where selected groups of deregulated genes indeed indicate high level of cell-type specificity, such as for instance the elevated reactive oxygen species (ROS) upon loss of *Rptor*. Critical insight into cell-type specificity of mTORC1 signaling also comes from recent studies that analyzed gene expression at single cell level in samples of patients suffering from FCD (Baldassari et al. 2025; Bizzotto et al. 2025). Intriguingly, only a small fraction (less than 10%) of mutated cells - with activating mTOR mutations - in the analyzed FCD samples exhibited the characteristic aberrant cytomegalic features. Thus the sensitivity to over-activation of mTORC1 in FCD exhibits major cell-type specificity (Baldassari et al. 2025). Even more important, cell-type specific transcriptional dysregulation was not only observed in mutated cells but also in cells that did not carry the mutation in the mosaic FCD tissue. Thus cell-autonomous and non-cell-autonomous, cell-type specific features upon mTORC1 over-activation may underlie the etiology of clinical symptoms, such as refractory epilepsy in FCD patients (Baldassari et al. 2025; Bizzotto et al. 2025). It will be important in future studies to investigate how cell-type specificity of mTORC1 signaling, or the loss thereof, is established and regulated throughout development at the cellular level; and how posttranscriptional mechanisms further fine tune cell-type specificity of intracellular signaling in health and disease.

## METHODS

### Maintenance and breeding of mice

All mouse colonies were kept according to protocols approved by institutional animal care and use committee, institutional ethics committee and the preclinical core facility (PCF) at the Institute of Science and Technology of Austria (ISTA). Experiments were performed under a license approved by the Austrian Federal Ministry of Women, Science and Research in accordance with the Austrian and EU animal laws (license numbers: BMWF-66.018/0007-II/3b/2012, BMWFW-66.018/0006-WF/V/3b/2017, and GZ 2025-0.597.515). All mice used in this project showed specific pathogen free status according to FELASA recommendations (Mahler et al. 2014). Animals were maintained and bred in the Preclinical Core Facility (PCF) of ISTA in accordance of EU regulations: room temperature 21 ± 1°C; relative humidity 40%–55%; photoperiod 12L:12D. Food (V1126, Ssniff Spezialdiäten GmbH, Soest, Germany) and tap water available *ad libitum*.

Mouse lines with MADM cassettes inserted on Chr. 4 and Chr. 11: MADM-4-GT (Contreras et al. 2021), MADM-4-TG (Contreras et al. 2021), MADM-11-GT (Hippenmeyer et al. 2010) (JAX stock # 013749), MADM-11-TG (Hippenmeyer et al. 2010) (JAX stock # 013751); *Bax*^flox^, *Bak*^flox^ (Takeuchi et al. 2005) (JAX stock #006329); *Emx1*-*Cre* (Gorski et al. 2002) (JAX stock #005628); *Emx1-CreER* (Kessaris et al. 2006) (JAX stock #027784); *Mtor* (Risson et al. 2009) (JAX stock #011009); *p53* KO (Jacks et al. 1994) (JAX stock #008462); and *Rptor^flox^* (Sengupta et al. 2010) (JAX stock #013188) have been described previously. We have not observed any influence of sex on the results in our study, and all experiments and analyses utilized animals independent of their sex. All efforts were made to minimize the number of animals used following the 3R principles. All MADM-induced analyses in animals were carried out in a mixed CD1-C57BL/6J genetic background with the exception of MADM-4 in a CD1-C57BL/6N background. Animals from forward (matUPD in red; patUPD in green) and/or reverse (matUPD in green and patUPD in red) crossing schemes were used for analysis and data acquisition.

### Generation of MADM recombinants and experimental mice

To generate MADM genetic mosaic mice for *Rptor* gene, we followed previously established protocols (Amberg and Hippenmeyer 2021; Contreras et al. 2021). In brief, *Rptor* mutant allele was recombined onto chromosome 11 containing the MADM-TG cassette to generate *MADM-11^TG/TG,Rptor^* stocks. Next, *MADM-11^TG/TG,Rptor^*were crossed with *MADM-11^GT/GT^*;*Emx1^Cre/+^* or *MADM-11^GT/GT^*;*Emx1-CreER^+/-^* to generate Control-MADM (*MADM-11^GT/TG^*;*Emx1^Cre/+^* or *MADM-11^GT/TG^*;*Emx1-CreER^+/-^*) and *Rptor*-MADM (*MADM-11^GT/TG,Rptor^*;*Emx1^Cre/+^*or *MADM-11^GT/TG,Rptor^*;*Emx1-CreER^+/-^*) mice (Figure S2A). Upon *Cre* recombinase-mediated interchromosomal recombination for reconstitution of the fluorescent MADM markers, all (red tdT^+^, green GFP^+^ and yellow tdT^+^/GFP^+^) cells in Control-MADM were wild-type (*Rptor^+/+^*) whereas in *Rptor*-MADM red tdT^+^ cells were wild-type (*Rptor^+/+^*), green GFP^+^ cells homozygous mutant (*Rptor^-/-^*) and yellow tdT^+^/GFP^+^ cells heterozygous (*Rptor^+/-^*) in an otherwise unlabeled *Rptor^+/-^*environment (Figures S1 and S2). In order to generate the *Rptor* conditional knock-out *(MADM-11^GT,Rptor/TG,Rptor;Emx1Cre/+^), MADM-11^TG/TG,Rptor^* was crossed with MADM-11*^GT/GT^*, Rptor;Emx*^1Cre/+^* (Figure S2B).

*Mtor* gene is located on chromosome 4. Briefly, *Mtor* mutant allele was recombined onto chromosome 4 containing the MADM-TG cassette to generate *MADM-4^TG/TG,Mtor^* stocks. Next, *MADM-4^TG/TG,Mtor^*mice were crossed with *MADM-4^GT/GT^*;*Emx1^Cre/+^*to generate Control-MADM (*MADM-4^GT/TG^*;*Emx1^Cre/+^*) and *Mtor*-MADM (*MADM-4^GT/TG,Mtor^*;*Emx1^Cre/+^*) mice.

For the epistasis experiments, we incorporated mutations of *Trp53* or *Bax/Bak* genes into our MADM paradigms. *Trp53* is located on chromosome 11 like *Rptor*. *Trp53* mutant allele was recombined onto chromosome 11 containing the MADM-TG cassette and the *Rptor* mutant allele (*MADM-11^TG/TG, Rptor^*) generating the parental MADM mice *MADM-11^TG/TG,Rptor,Trp53^.* Next, *MADM-11^TG/TG,Rptor,Trp53^*were crossed with *MADM-11^GT/GT^*;*Emx1^Cre/+^* to generate *Rptor-Trp53*-MADM (*MADM-11^GT/TG,Rptor,Trp53^*;*Emx1^Cre/+^*) mice (Figure S9A).

*Bax* and *Bak* genes are located on two different chromosomes (7 and 17 respectively). In order to generate the parental MADM mice for the *Rptor-Bax*-MADM, *Bax^flox/+^*;*Bak^Delta/+^* animals were crossed with *MADM-11^TG/TG,Rptor^* generating *MADM-11^TG/TG,Rptor^;Bax^flox/+^*;*Bak^Delta/+^*. *Bax^flox/+^*;*Bak^Delta/+^* animals were breed with *MADM-11^GT/GT^*;*Emx1^Cre/+^* to obtain the parental mice *MADM-11^GT/GT^;Bax^flox/+^*;*Bak^Delta/+^;Emx1^Cre/+^*. By crossing both parental mice: *MADM-11^TG/TG,Rptor^;Bax^flox/+^*;*Bak^Delta/+^*and *MADM-11^GT/GT^;Bax^flox/+^*;*Bak^Delta/+^;Emx1^Cre/+^*, the ultimate experimental *MADM-11^GT/TG,Rptor^;Bax^flox/flox^;Bak^Delta/Delta^;Emx1^Cre/+^*mice were generated (Figure S9B).

### Isolation of tissue and immunohistochemistry

Postnatal mice were deeply anesthetized by intraperitoneal (IP) injection of a ketamine/xylazine solution (65 mg, 13 mg/kg body weight, respectively), and unresponsiveness was confirmed through pinching in the paw. The diaphragm of the mouse was opened from the abdominal side to expose the heart. Cardiac perfusion was performed with ice-cold PBS followed immediately by 4% PFA prepared in PB buffer (Sigma-Aldrich). Brains were removed and further fixed in 4% PFA o/n to ensure complete fixation. For embryos, pregnant female mice were sacrificed by cervical dislocation and embryonic heads were fixed by immersion in 4% PFA ice-cold o/n. All brains were cryopreserved with 30% sucrose (Sigma-Aldrich) solution in PBS for approximately 48 hours. Brains were embedded in Tissue-Tek O.C.T. (Sakura). For adult time points, 40µm coronal sections were collected in 24 multi-well dishes (Greiner Bio-one) and stored at −20°C in antifreeze solution (30% v/v ethyleneglycol, 30% v/v glycerol, 10% v/v 0.244M PO_4_ buffer) until used.

Embryonic brains were sectioned at 20µm and directly mounted onto Superfrost glass-slides (Thermo Fisher Scientific) for storage at −20 ° C. For immunohistochemistry, adult brain sections were mounted, followed by 3 wash steps (5min) with PBS. All tissue sections were blocked for 30 minutes in a blocking buffer solution of 5% normal donkey serum (Thermo Fisher Scientific), 0.3% Trition X-100 in PBS. Primary antibodies were prepared in blocking buffer and incubated o/n at 4°C. Sections were washed 3 times for 5 minutes each with PBS-T (0.3% Triton X-100 in PBS) and incubated with corresponding secondary antibody diluted in PBS-T for at least 1 hour. Sections were washed twice with PBS-T and once with PBS. Slices were incubated in 2.5% DAPI for nuclear staining (Thermo Fisher Scientific). For the analysis of axonal projections, sections were at 100μm and immunohistochemistry was performed in free-floating following the same process as described above. Sections were embedded in mounting medium containing 1,4-diazabicyclooctane (DABCO; Roth) and Mowiol 4-88 (Roth) and stored at 4°C.

### MADM clonal labeling experiments

MADM clonal analysis was performed as previously described (Beattie et al. 2020). Briefly, pregnant females carrying E11 or E12 *Rptor*-MADM;*Emx1-CreER^+/-^*embryos were IP injected with 100µl of 20mg/ml Tamoxifen (TM) to induce clonal MADM labeling. For the analysis, E13 or E16 embryos; P0 or P21 pups were fixed and processed as specified above for immunochemistry.

### EdU labeling experiments

Cycling RGPs were assessed by EdU incorporation. Experiments were based on the use of the Click-iT Alexa Fluor 647 imaging kit (Thermo Fisher). Reagents were reconstituted according to the user manual. Intraperitoneal (IP) EdU injections were performed at E17, one hour before fixing (1mg/ml EdU stock solution; 30-40ml per mouse) to label S-phase cells. Tissue collection and immunohistochemistry was performed as described above. The Click-iT imaging kit was used according to the instruction manual to visualize the EdU signal before the DAPI staining was performed.

### Imaging and analysis in MADM-labeled brains

Sections were imaged using either an inverted LSM800 or LSM880 with airy scan confocal microscope (Zeiss) and processed using Zeiss Zen Blue software. Tiled images were taken for at least three brain sections per animal. MADM-labeled cells were manually counted based on respective marker expression. Quantification of EdU, PH3 and Caspase-3 labeling was performed using a 40x oil objective.

### Volume quantification of axonal projections

In order to trace the axonal projections of cortical neurons we generated 100µm thick brain sections by using a cryostat. We used the same immunostaining protocol previously described in this manuscript but on free-floating. After anti-GFP, anti-tdTomato and DAPI staining we mounted the entire brain section onto Superfrost glass-slides and embedded them in DABCO-Mowiol mounting medium. The sections were imaged in a Nikon Ti2E inverted microscope (Yokogawa CSU-W1 dual-disk spinning disk unit) and an Apo λ 20x air 0.75/0.17-0.25mm objective. For the analysis, we used Fiji to crop the areas of interest, split the channels and export the TIFF files. Then, using Ilastik 1.4 software, we used machine learning based pixel classification and segmentation, converting our images into binary images keeping the z-stack intact (feature selection 0.3-1 sigma). With Fiji, we imported the generated files and subtracted the green channel from the red channel and the red channel from the green channel at each z-stack to eliminate the signal from GFP^+^tdTomato^+^ (yellow) cells. We selected a ROI around the track and cleared the outside to remove signal from cell bodies or other regions. Finally, we calculated the total volume of the projections of each subtracted channel by the tool MorphoLibJ (Analyze regions 3D). The G/R ratio was calculated for each hemisphere and averaged in each section for all relevant genotypes.

### Preparation of single cell suspension and FACS for RNA sequencing and proteomics

Single cell suspensions of MADM labelled cells were prepared as described (Laukoter et al. 2020a; Amberg et al. 2024). Experimental animals at E13, E16 or P0 were sacrificed, brains extracted and cortex dissected. Single cell suspensions were obtained by using Papain containing L-cysteine and EDTA (vial 2, Worthington), DNase I (vial 3, Worthington), Ovomucoid protease inhibitor (vial 4, Worthington), EBSS (Thermo Fisher Scientific), DMEM/F12 (Thermo Fisher Scientific), FBS (Thermo Fisher Scientific) and HS (Thermo Fisher Scientific). All vials from Worthington kit were reconstituted according to the manufacturer’s instructions using EBSS. The dissected brain areas were directly placed into Papain-DNase solution (20 units/ml papain and 1000 units DNase). Enzymatic digestion was performed for 15-30min at 37°C in a shaking water bath. Next, solution 2 (EBSS containing 0.67mg Ovomucoid protease inhibitor and 166.7 U/ml DNase I) was added. The whole suspension was thoroughly mixed and centrifuged for 5min at 1000rpm at RT. Supernatant was removed and cell pellet was resuspended in solution 2. Trituration with p1000 pipette was performed to mechanically disaggregate any remaining tissue parts. DMEM/F12 was added to the cell suspension as a washing solution, followed by a centrifugation step of 5min with 1500rpm at RT. Cells were resuspended in media (DMEM/F12 containing 10% FBS and 10% HS) and kept on ice until sorted. Right before sorting, cell suspension was filtered using a 40mm cell strainer. FACS was performed on a BD FACS Aria III or Sony SH800SFP using 100 nozzle, and keeping sample and collection devices (1.5ml tubes or 96-well plate) at 4°C. Duplet exclusion was performed to ensure the sorting of true single cells. For single cell analysis, individual GFP^+^ cells for each condition were sorted.

### Sample preparation for proteomics

After obtaining each cell suspension in Eppendorf tubes, samples were immediately frozen by liquid nitrogen immersion. First, we pooled the samples to have 10000-20000 cells per each replica. To pool cell samples, the samples were kept in dry ice, then normal ice, lysate buffer was added to the first sample, the liquid dragged to the second sample and so on. 10 min at 95°C 1000rpm.

Frozen cell pellets were placed on dry ice, and processed with the iST-NHS kit (PreOmics GmbH) according to the manufacturer’s protocol with the following modifications: 100 µL LYSE-NHS was used per sample. Digestion was stopped after 3h. Three mixed development stage-specific internal reference samples were created by mixing 1/10th (E13 and E16) or 1/7th (P0) of each sample. Samples were then labelled with TMT-10plex (ThermoFisher Scientific), using 62 µL out of 150 µL per sample. After 1h, the reaction was quenched with 15 µL hydroxylamine, then 100 µL STOP was added. Individual samples were then combined into 6 combined TMT samples. Cleaned-up samples were then dried overnight, re-dissolved in 300 µL 0.1% TFA with 5 min sonication, then fractionated into 9 fractions with the Pierce High pH Reversed-Phase Peptide Fractionation Kit (ThermoFisher Scientific). The fractions, as well as the flow through and wash, were then dried, re-dissolved in 35 µL LC-LOAD and analysed by Mass Spectrometry. All LC-MS/MS runs were acquired on a Q Exactive HF (ThermoFisher Scientific) coupled with an Ultimate 3000 RSLC_Nano nano-HPLC (ThermoFisher Scientific). Samples were pre-concentrated over an Acclaim PepMap trap column (5 µm C18-coated particles, 0.5 cm * 300 µm ID, ThermoFisher Scientific P/N 160454), then separated on an EasySpray PepMap RSLC column (2 µm C18-coated particles, 50 cm * 75 µm ID, ThermoFisher Scientific P/N ES903) and eluted over the following 60 min gradient: solvent A, MS-grade H₂O + 0.1% formic acid; solvent B, 80% acetonitrile in H₂O + 0.08% formic acid; constant 300 nL/min flow; B percentage: 5 min, 2%; 45 min, 31%; 65 min, 44%, followed immediately by a 5 min plateau at 90%. Mass spectra were acquired in positive mode with the following Data Dependent Acquisition (DDA) method: chromatographic peak width FWHM = 20 s, MS1 parameters: 1 microscan, 120,000 resolution, AGC target 3e6, 50 ms maximum IT, 380 to 1500 m/z; up to 20 MS2s per cycle. DDA MS2 parameters: Centroid mode, 1 microscan, 60,000 resolution, AGC target 1e5, 100 ms maximum IT, 0.7 m/z isolation window (no offset), 380 to 1500 m/z, NCE 32, excluding charges 1+, 8+ and higher or unassigned, 10s dynamic exclusion.

### ROS detection and quantification

Single cell suspension of MADM labelled cells was prepared as described above with small modifications to include a staining step. Experimental animals at P0 or P2 were sacrificed by decapitation, brains extracted and cortex dissected. Single cell suspensions were obtained by using Papain containing L-cysteine and EDTA (vial 2, Worthington), DNase I (vial 3, Worthington), Ovomucoid protease inhibitor (vial 4, Worthington), EBSS (Thermo Fisher Scientific), DMEM/F12 (Thermo Fisher Scientific), FBS (Thermo Fisher Scientific) and HS (Thermo Fisher Scientific). All vials from Worthington kit were reconstituted according to the manufacturer’s instructions using EBSS. The dissected brain areas were directly placed into Papain-DNase solution (20 units/ml papain and 1000 units DNase). Enzymatic digestion was performed for 20min at 37°C in a shaking water bath. Next, solution 2 (EBSS containing 0.67mg Ovomucoid protease inhibitor and 166.7 U/ml DNase I) was added. The whole suspension was thoroughly mixed and centrifuged for 5min at 1000rpm at RT. Supernatant was removed and the cell pellet was resuspended in solution 2. Trituration with p1000 pipette was performed to mechanically disaggregate any remaining tissue parts. After centrifugation for 5 min at 100rpm, CellROX™ antibody was added in DMEM (1:500) and the sample incubated for 30min in a water bath at 37°C at 150rpm. Filtered PBS 1x was added to the cell suspension as a washing solution 3 times, followed by a centrifugation step of 5min with 1500rpm at RT. Cells were resuspended in media (DMEM/F12 containing 10% FBS and 10% HS) and kept on ice until sorted. Right before sorting, cell suspension was filtered using a 40mm cell strainer. FACS was performed on a Sony SH800SFP using 100µm chip and keeping sample and collection devices (1.5ml tubes) at 4°C. Duplet exclusion was performed to ensure the sorting of true single cells. Unlabeled samples and ROS^+^ only samples were used as negative and positive controls respectively. Individual GFP^+^ROS^+^ and GFP^+^ROS^-^ cells for each condition were sorted for subsequent analysis by bulk RNA-seq. The obtained data from the Flow cytometer was analyzed by Flow Cytometry Analysis Software FlowJo v10.8.1.

### RNA extraction and cDNA library preparation of MADM samples for RNA sequencing

Bulk RNA-seq was performed as described earlier (Laukoter et al. 2020b). In brief, RNA was isolated from sorted cells, using Trizol LS (Thermo Fisher Scientific) and dissolved in RNase-free H_2_O. RNA quality was analyzed using Bioanalyzer RNA 6000 Pico kit (Agilent) following the manufacturer’s instructions. High quality RNA samples were used for SMART-Seq2 (Picelli et al. 2014) or SMART-Seq3 (Hagemann-Jensen et al. 2020) library production and sequencing (Illumina platforms) was performed by the Next Generation Sequencing Facility at Vienna BioCenter Core Facilities (VBCF), member of the Vienna BioCenter (VBC), Austria (https://www.viennabiocenter.org/vbcf/).

For single cell RNA sequencing (scRNA-seq) libraries were generated using the Chromium iX Controller and the Next GEM Single Cell 3’ Reagent Kit (v3.1, 10x Genomics) according to the manufacturer’s instructions. Single cell suspensions were incubated with DNA-labeled cholesterol-modified oligos (Sigma) at 4°C. Following incubation, cell suspensions were washed two times with PBS containing BSA. After the final wash, cell suspensions were resuspended in PBS containing BSA and Zombie NIR fixable viability dye (Biolegend) for discrimination between live and dead cells. Next, viable GFP^+^ cells were sort-purified on a Sony SH800 cell sorter and cells obtained from E18 Control-MADM and E18 *Rptor*-MADM, as well as P1 Control-MADM and P1 *Rptor*-MADM. Samples were pooled for processing as a single sample according to the manufacturer’s protocol, with the cholesterol-linked barcodes enabling sample-specific demultiplexing of the sequencing data. Libraries were sequenced by the Biomedical Sequencing Facility at the CeMM Research Center for Molecular Medicine of the Austrian Academy of Sciences, using the Illumina NovaSeq 6000 platform. Raw sequencing data was pre-processed and demultiplexed using Cell Ranger (v7.0, 10x Genomics).

### Processing and statistical analysis of bulk RNA-seq data to analyze developmental time course of Control-MADM, *Rptor*-MADM, and cKO-*Rptor*-MADM

Reads were aligned to: GRCm38.p5b with Gencode M16 annotation (https://www.gencodegenes.org) using STAR aligner. For downstream analysis, count tables from STAR were used for analysis with R v4.3.2. We performed principal component analysis (PCA) of the top 500 variable genes in all analyzed samples and removed a single sample based on its position in the PCA plot (PCA of the final set of samples is shown in Figure S3A). We retained 2-7 samples per replicate, which showed a unique alignment rate between 69% - 79% resulting in 3.9M – 19.7M reads for downstream analyses. Figure S3B: The conditional deletion allele analyzed here removed *Rptor* exon 6 (Sengupta et al. 2010). Therefore, we quantified reads in this region (chr11:119756238-119756414, GRCm38/mm10) to test for efficient recombination. Read counts were then incorporated into the count tables for downstream analyses. Figure shows the normalized read counts, determined by DESeq’s counts function. Differential gene expression was calculated using DESeq2 separately for each developmental time point using contrasts to compare pairwise comparisons of genotypes. Only genes with a mean read count >10 in all samples were used for further analysis. Figure 3B, D, S3C: Number and overlap statistics of differentially expressed genes (DEG) from comparison of cKO-*Rptor*-MADM/Control-MADM or *Rptor*-MADM/Control-MADM with an adjusted p-value < 0.05 and a log2 fold-change > 0 (up, higher in respective *Rptor* deletion cells compared to Control-MADM cells) or < 0 (down, lower in respective *Rptor* deletion cells compared to Control-MADM cells). Figure S3D: We created a set of 376 manually curated GO terms, that are potentially connected to *Mtor/Rptor* function, to determine GO term enrichment in DEGs with the enricher function of the clusterProfiler package. We focused on GO terms with an uncorrected p-value < 0.05 and calculated a score for each GO term as the negative log10 of the uncorrected p-value. GO terms were grouped via presence of terms axon (Axon), cell cycle (Cell cycle), mitoch (Mitochondria), TOR (TOR), apop (Apoptosis), reactive_oxygen (ROS), metabol (Metabolism) or cytokine (Cytokine) in the GO term description. The highest score for each GO term group and sample (as indicated in on the x-axis) was used to draw the heatmap. Scores were cut at 2.5 for better visualization. Figure 3F, S3E: For STRINGdb analysis we focused only on *Rptor*-MADM/Control-MADM comparison and used DEGs unique to one developmental time point (212 DEGs). A protein-protein-interaction network was constructed using the STRING database (v12) (Szklarczyk et al. 2023) and the stringdb R package with standard settings and interaction score cutoff: 400. Networks were visualized using Cytoscape and RCy3 R package (Gustavsen et al. 2019). We used the largest connected network (79 genes) for further analysis. Genes were assigned to GO term groups via GO term descriptions.

### Processing and statistical analysis of proteomic data

Raw files were searched in MaxQuant against a fasta proteome downloaded from UniProtKB. Fixed cysteine modification was set to +C6H11NO; variable modifications were set to Acetyl (protein N-term), Oxidation (methionine), Gln->pyroGlu and Deamidation (NQ). Match between runs was turned on. All data was filtered at 1% FDR.

MaxQuant’s output was post-processed using in-house scripts, starting from the evidence.txt output file. TMT reporter intensities were corrected for TMT purity factors, scaled to MS1 intensities, normalised using the Levenberg Marquardt procedure to minimize sample-to-sample differences, then batch corrected using the Internal Reference Scaling procedure using reference channels. Protein groups were inferred and quantified using an in-house MaxLFQ-like algorithm. Protein groups were re-normalised and tested using limma. GO terms enrichment analysis was based on topGO.

For Figure 3C, E, S3F number and overlap statistics of genes were connected to differentially expressed proteins (DEP) from comparison of cKO-*Rptor*-MADM/Control-MADM or *Rptor*-MADM/Control-MADM with an FDR < 1% and tagged “up” (higher in respective *Rptor* deletion cells compared to Control-MADM cells) or tagged “down” (lower in the respective *Rptor* deletion cells compared to Control-MADM). GO term enrichment was calculated and plotted as for DEGs (Figure S3G). Due to the large amounts of DEPs, we focused this analysis on genes connected to one or more GO terms (same as used for GO term enrichment analysis) (Figure 3G, S3H). Only DEPs specific to one developmental time point were used for the analysis (314 genes). A protein-protein-interaction network was constructed using the STRING database (v12) and the stringdb R package with standard settings and interaction score cutoff: 400. We used the largest connected network (285 genes) for further analysis. Networks were visualized using Cytoscape and RCy3 R package. Genes were assigned to GO term groups via GO term descriptions.

### Processing and statistical analysis of bulk RNA-Seq data to analyze cells with elevated ROS

Reads were aligned to GRCm39 with Gencode M27 annotation (https://www.gencodegenes.org) using STAR aligner. For downstream analysis, count tables from STAR were used for analysis with R v4.3.2. We removed 4 samples due to low alignment rate <50%. We performed principal component analysis (PCA) of the top 500 variable genes in all analyzed samples and removed a single sample based on its position in the PCA plot. We retained 3-5 replicates per sample, which showed a unique alignment rate between 50% - 71% resulting in 1.7M – 3.9M uniquely aligned reads for downstream analyses. Note that the sequencing depth was not sufficient to analyze expression of the deleted *Rptor* exon 6. DEG statistics were determined using DESeq2 using contrasts to compare genotypes. Marker genes for cortical cell types were identified using published scRNA-Seq data of P1 mice (GEO: GSM4635080). Note that several deep layer cell types were combined for this analysis: DL CPN, CThPN, SCPN, Layer 4, Layer 6b, NP. Marker genes as well as DEG statistics were used to perform gene set enrichment analysis using GSEA function from clusterProfiler. Enrichment scores were calculated as the negative log10 of the adjusted p-value. Only scores for deep-, and upper layer neurons are shown in Figure S9D.

### Analysis of scRNA-seq data

All analyses were performed with R v4.3.2 (unless indicated otherwise). Filtering and initial analysis of scRNA-seq data was performed using Seurat R package, following standard pipelines. In brief, parameters for filtering high quality cells were: nFeature_RNA > 200 & nFeature_RNA < 5500 & nCount_RNA < 20000 & percent.mt < 5 (Figure S6A and S6B). Partial cell cycle regression was performed using Seurat’s CellCycleScoring function and cell cycle genes from: https://github.com/hbc/tinyatlas/tree/master/cell_cycle/Mus_musculus.csv. Label transfer of cell types from a published reference (P1 cortex GEO: GSM4635080, (Di Bella et al. 2021) was performed using FindTransferAnchors with dims = 1:20 and TransferData with dims = 1:15 (Figure S6C and S6E). UMAP and clustering analysis was performed using RunUMAP with dims = 1:30, FindNeighbors with dims = 1:30 and FindClusters with resolution = 1 (Figure S6C-F). Post mitotic neurons were annotated as clusters that expressed established marker genes (*Dcx*, *Neurod1*, Figure S6G-S6H, Figure 6B), identified as migrating neurons by the reference cell type annotation and were assigned to a single genotype (*Rptor*-MADM, Control-MADM) during the demultiplexing procedure. *Satb2* and *Ctip2* (*Bcl11b*) expressing cell types were identified using AddModuleScore and the following gene sets: *Satb2*: *Satb2*, *Cux1*, *Ppfia2*, *Frmd4a, Cux2, Robo2, Gria2, Hs6st2, Clstn2, Rbfox1, Ptprz1, Gpm6a, Ptn, Ttc28, Osbpl6; Ctip2: Bcl11b, Foxp2, Fezf2, Pcp4, Rprm, Synpr, Zfpm2, Rnd2* (Figure 6C and S6I). We validated this approach by determining DEGs between *Satb2* and *Ctip2* cells and plotting an expression heatmap for genes with adjusted p < 10^-20^ (Figure S6I) as well as plotting expression of selected marker genes in pseudobulk data created with AggregateExpressionCells (Figure 6C). For each developmental time point (E18, P1) and cell type (*Satb2*, *Ctip2*) we determined differential expression statistics for *Rptor*-MADM/Control-MADM using FindMarkers with test.use = “MAST” and logfc.threshold = 0.01 parameters. These statistics were used for Integrative Differential expression and gene set Enrichment Analysis (iDEA, R v4.4.1) and the iDEA results were used for downstream analyses. GO terms were ranked by adjusted p-value and the top 50 GO terms used for further analyses. GO terms were grouped using simplifyGO function from simplifyEnrichment package. GO groups were assigned manually based on clusters of GO terms identified by simplifyGO. Enrichment score was calculated as the negative log10 of the adjusted p-value, cut at 2 and was used for plotting heatmaps (Figure 6E-6F and S7). In an attempt to give a concise overview of the relatively long lists of GO term descriptions, we calculated the frequency of each word in the GO term descriptions for each GO term cluster (i.e. fraction of GO term descriptions containing the respective word). For the figures, subsets of GO term groups, clusters and word frequencies were chosen arbitrarily and displayed. We focused on E18 and analyzed GO terms that were significantly enriched (score > 1.5) in only one comparison (Figure 6F). Posterior inclusion probabilities (PIPs) from all genes in the respective GO terms were extracted and the maximum PIP for each gene was used for further analyses. Top 10 genes (ranked by PIP value) were identified for *Satb2* and *Ctip2* cell types and PIP values plotted.

### Quantitative analysis of MADM-labeled brains

Sections were imaged using an inverted LSM800 confocal microscope (Zeiss) or epifluorescence microscopy (Olympus VS120 Slide scanner) and processed using Zeiss Zen Blue software or OlyVIA software respectively. Confocal images were imported into Photoshop software (Adobe) or Fiji ImageJ software and MADM-labeled cells were manually counted based on respective marker expression (Beattie et al. 2017). Statistical analysis was performed in Graphpad Prism 7.0. All data used for quantification of MADM-labeled tissue is compiled in Table S1.

## ACKNOWLEDGMENTS

We thank A. Heger (IST Austria Preclinical Facility), A. Sommer (VBCF GmbH, NGS Unit), and A. Nicolas (IST Austria Lab Support Facility / Mass Spectrometry Facility) for technical support; K. Ferencak, I. Aykara, P. Hirschfeld, E. Fisher, S. Laukoter, L. Andersen for initial experiments and/or assistance; and all members of the Hippenmeyer lab for discussion. This research was supported by the Scientific Service Units (SSU) of IST Austria through resources provided by the Imaging and Optics- (IOF), Lab Support-(LSF) and Preclinical Facilities (PCF). R.B. received support from FWF Meitner-Programm (M 2416). This work was also supported by IST Austria institutional funds; the People Programme (Marie Curie Actions) of the European Union’s Seventh Framework Programme (FP7/2007-2013) under REA grant agreement No 618444 to S.H., and the European Research Council (ERC) under the European Union’s Horizon 2020 research and innovation programme (grant agreement No 725780 LinPro) to S.H.

## Author Contributions

S.H. conceived the research. S.H., R.B., A.V., and F.M.P. designed all experiments and interpreted the data. R.B., A.V., F.M.P., C.S., T.K., M.S. and M.F. performed all the experiments. C.B. contributed expertise for the scRNA-seq experiments. F.M.P. performed all computational and bioinformatics analysis with inputs from A.V., R.B., O.A.M. and S.H. T.R. was instrumental in the generation of MADM mouse models. S.H. and A.V. wrote the manuscript with inputs from R.B. and F.M.P. All authors edited and proofread the manuscript.

## Materials Availability

All published reagents and mouse lines will be shared upon request within the limits of the respective material transfer agreements.

## Data and Code Availability

All data generated and analyzed in this study are included in the paper and/or supplementary materials. Raw sequencing data will be submitted to Gene Expression Omnibus (GEO).

## SUPPLEMENTARY FIGURE LEGENDS

**Figure S1.**
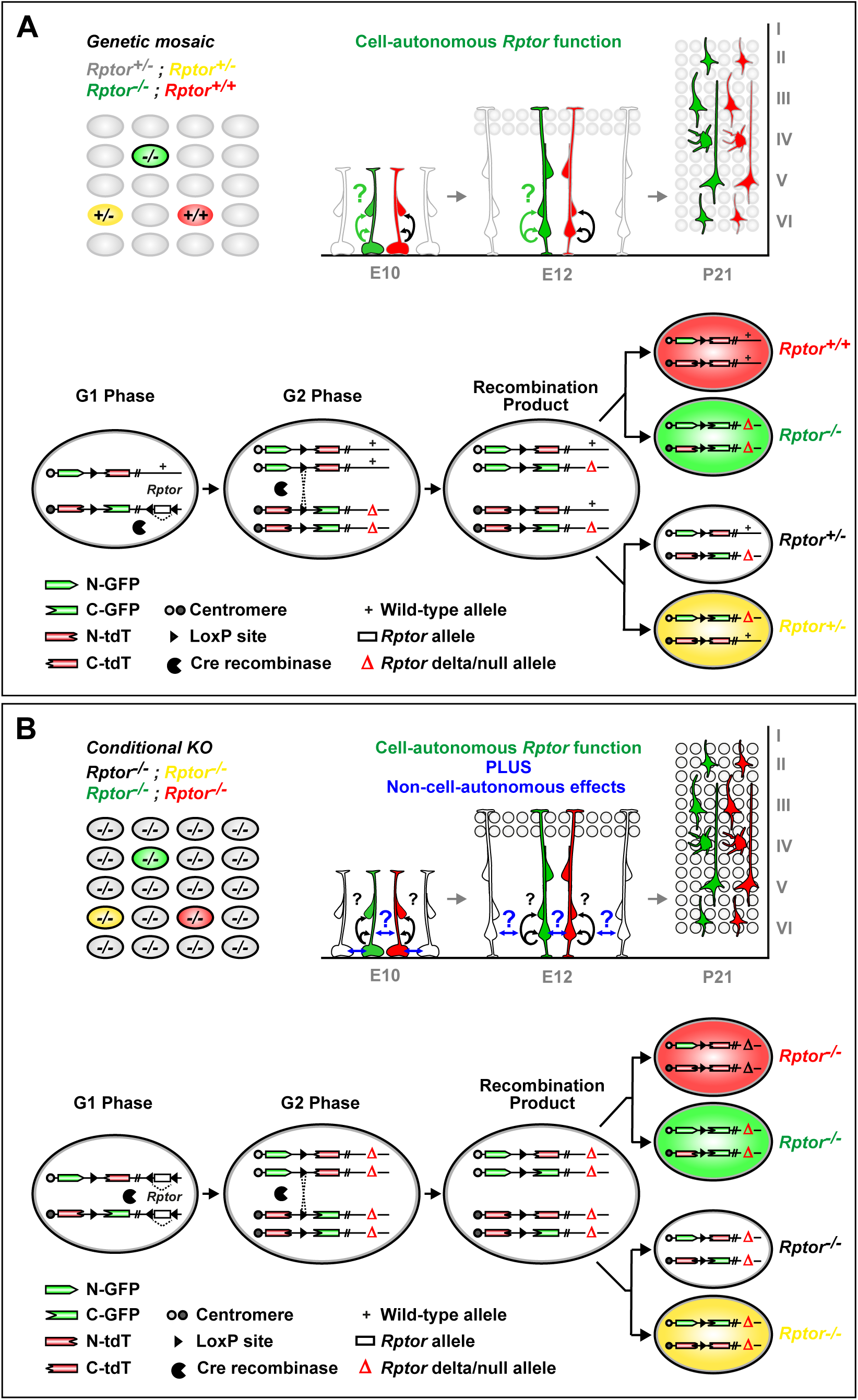
Experimental MADM Principle for Sparse and Global Gene Knockout with Double Marker Labeling at Single Cell Resolution. (A) (Top) Illustration of sparse genetic mosaic MADM paradigm with uniquely labeled homozygous *Rptor^-/-^* mutant (green), *Rptor^+/+^* control (red) and *Rptor^+/-^* (yellow) cells in an otherwise normal (provided no gene dosage sensitivity) heterozygous unlabeled background. The sparse mosaic (using constitutive Cre) or clonal (using TM-inducible CreER) MADM condition permits assessing the cell-autonomous *Rptor* gene function during cortical stem cell lineage progression in cortical development. (Bottom) Schematic representation of MADM principle to generate sparse genetic mosaics. The *Rptor-flox* allele was introduced distal to the TG-MADM cassette via meiotic recombination [for details on introducing mutant alleles into the MADM system see also (Hippenmeyer et al. 2010; Contreras et al. 2021)]. Cre recombinase mediates *cis*-recombination between the *LoxP* sites of the conditional *Rptor* allele (resulting in *Rptor-Δ*/null allele). Cre mediates as well interchromosomal *trans*-recombination between the MADM cassettes which, if happens in G2 phase, generates either one green *Rptor^-/-^* cell and one red *Rptor^+/+^* cell (upper branch), or one unlabeled *Rptor^+/-^* and one yellow *Rptor^+/-^*cell (lower branch). (B) (Top) Illustration of global/conditional (full) tissue-wide *Rptor* knockout with sparse MADM labeling. Green, red and yellow cells are all homozygous *Rptor^-/-^* mutant in a homozygous mutant unlabeled background. The sparse MADM labeling permits spatiotemporal phenotypic analysis of individual *Rptor^-/-^*mutant RGPs and their progeny at single cell resolution during cortical development. Note that any observed phenotype reflects the consequence of the loss of cell-autonomous *Rptor* gene function plus the influence and/or contribution of non-cell-autonomous and/or community effects. (Bottom) Schematic representation of MADM principle to generate conditional *Rptor* knockout with sparse MADM labeling. The *Rptor-flox* allele was individually introduced distal to both, the TG- and GT-MADM cassette via meiotic recombination. Cre recombinase mediates *cis*-recombination between the *LoxP* sites of the conditional *Rptor* alleles (resulting in *Rptor-Δ*/null alleles), as well as interchromosomal *trans*-recombination between the MADM cassettes, which generates either one green cell and one red cell (upper branch), or one unlabeled and one yellow cell (lower branch). All green, red and yellow cells are then homozygous *Rptor^-/-^* mutant in an unlabeled homozygous *Rptor^-/-^* mutant background. Note that green *Rptor^-/-^* mutant cells present an identical genotype in experimental paradigms in (A) and (B). Thus, any difference in the observed phenotype of green *Rptor^-/-^* mutant cells in the genetic mosaic (A) when compared to green *Rptor^-/-^* mutant cells in the conditional tissue-wide *Rptor* knockout (B) represents the contribution of non-cell-autonomous effects originating from the environment. Schematic is reused, adapted and modified from (Beattie et al. 2017), with permission.

**Figure S2.**
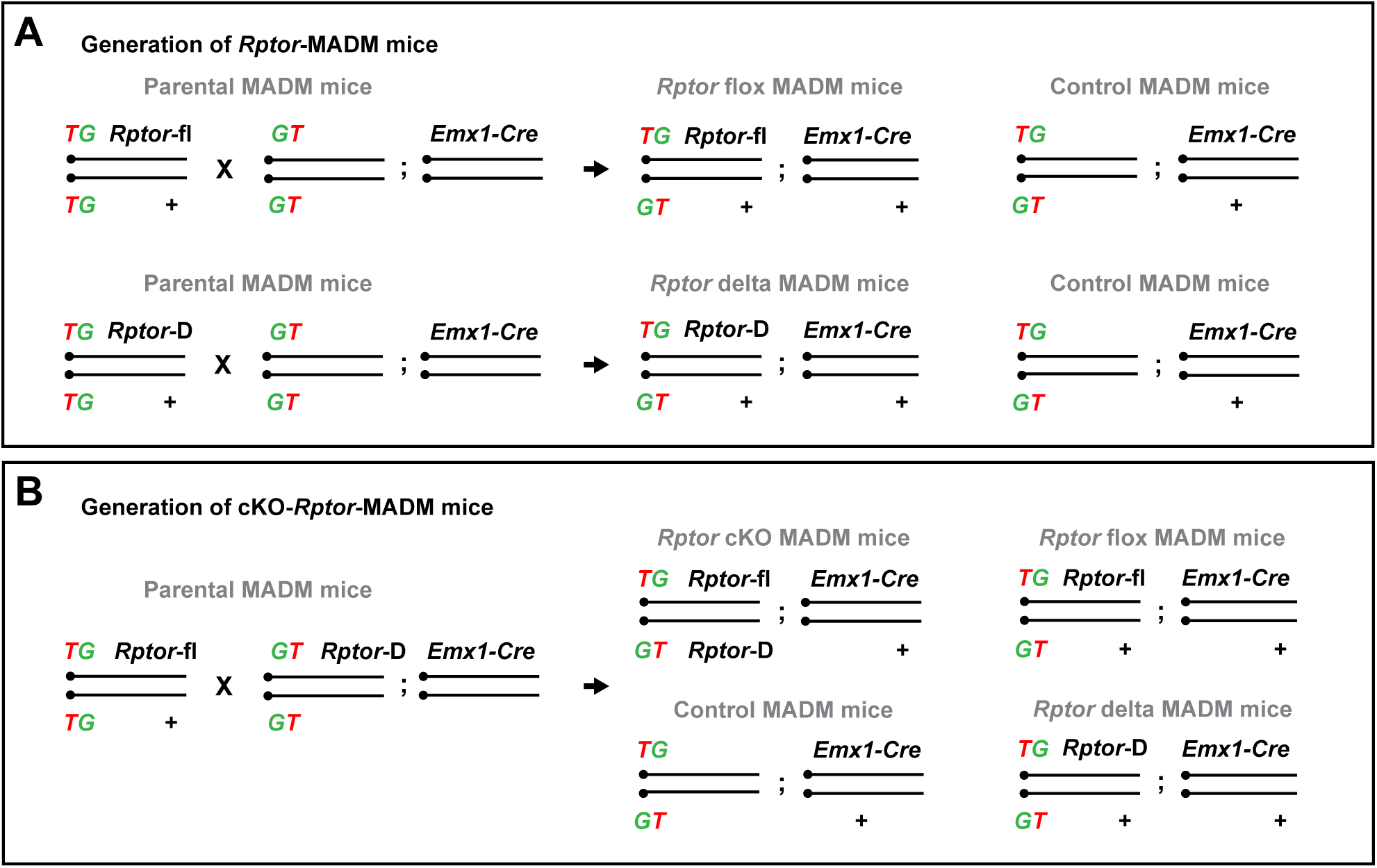
Breeding schemes for the generation of *Rptor*-MADM and cKO-*Rptor*-MADM experimental mice, related to Figures 2-6. (A) Breeding schemes for the generation of mosaic *Rptor*-MADM mice. *MADM-11^TG/TG,Rptor-flox(fl)^* or *MADM-11^TG/TG,Rptor-Delta(D)^* were crossed with *MADM-11^GT/GT^*;*Emx1^Cre/+^*to generate Control-MADM (*MADM-11^GT/TG^*;*Emx1^Cre/+^*) and *Rptor*-MADM (*MADM-11^GT/TG,Rptor-fl^*;*Emx1^Cre/+^*or *MADM-11^GT/TG,Rptor-D^*;*Emx1^Cre/+^*) mice. No phenotypic difference was observed between *MADM-11^GT/TG,Rptor-fl^*;*Emx1^Cre/+^*and *MADM-11^GT/TG,Rptor-D^*;*Emx1^Cre/+^* experimental animals. (B) Breeding schemes for the generation of cKO-*Rptor*-MADM mice. The parental mice *MADM-11^TG/TG,Rptor-fl^*and *MADM-11^GT/GT,Rptor-D^*;*Emx1^Cre/+^* were crossed to generate the cKO-*Rptor*-MADM (*MADM-11^GT,Rptor-D/TG,Rptor-fl^*;*Emx1^Cre/+^*), Control-MADM (*MADM-11^GT/TG^*;*Emx1^Cre/+^*) and *Rptor*-MADM (*MADM-11^GT/TG,Rptor-fl^*;*Emx1-Cre*^+/-^ or *MADM-11^GT/TG,Rptor-D^*;*Emx1-Cre*^+/-^).

**Figure S3.**
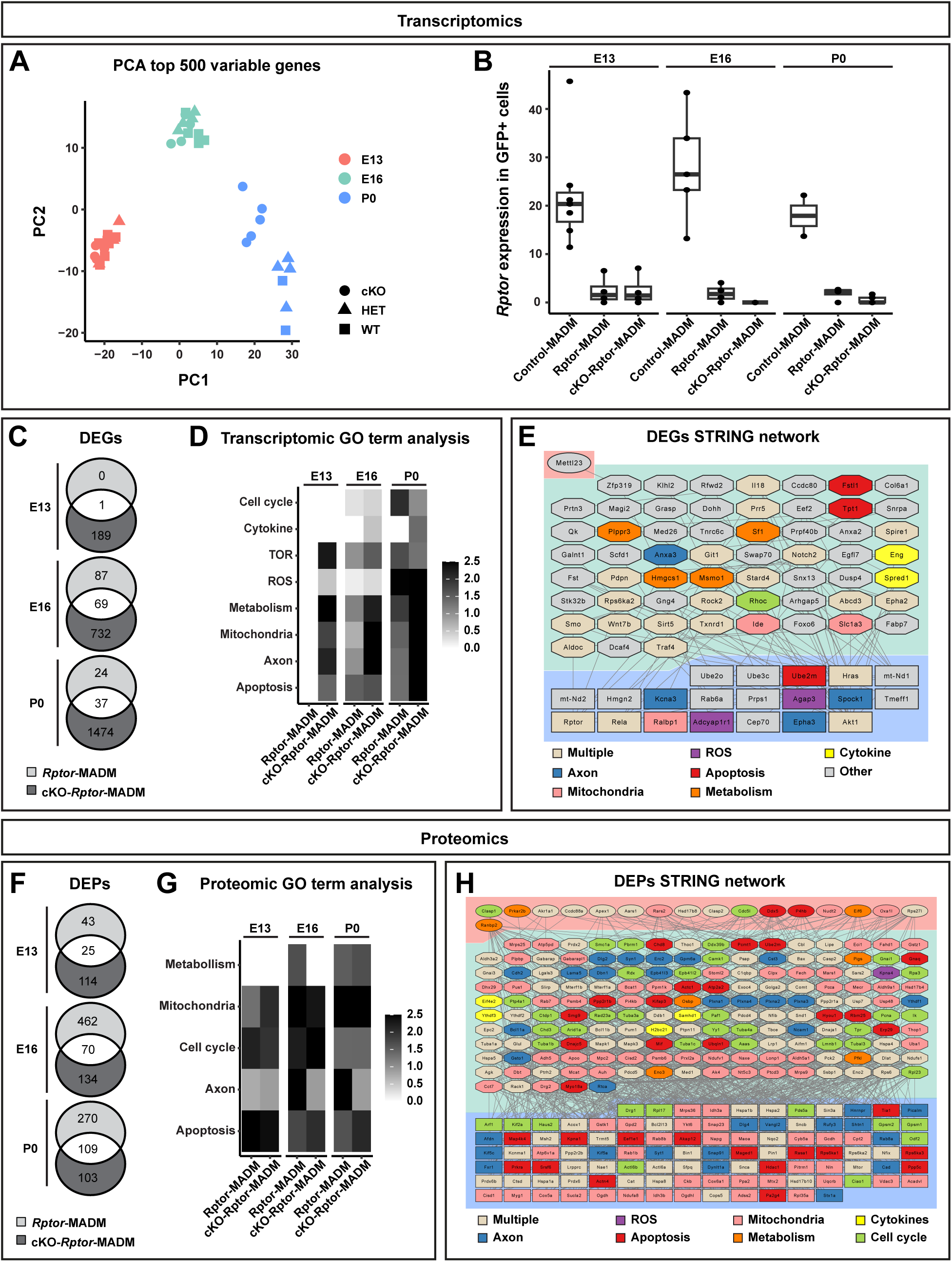
Analysis of transcriptomic and proteomic changes upon sparse and whole-tissue *Rptor* ablation, related to Figure 3. (A) PCA (top 500 variable genes) of all samples that were used for in-depth analyses. (B) Validation of *Rptor* deletion in MADM context. Normalized read counts in floxed *Rptor* exon 6, which was deleted in *Rptor*-MADM and cKO-*Rptor*-MADM. (C-H) Analysis of differential gene and protein expression upon sparse and whole-tissue *Rptor* ablation. (C and F) Overlap of differentially expressed genes (C, DEGs, adjusted p-value < 0.05 DESeq2) or genes connected to differentially expressed proteins (F, DEPs, FDR < 1% t-test) from comparison of *Rptor*-MADM/Control-MADM or cKO-*Rptor*-MADM/Control-MADM at different developmental time points (E13, E16, P0). (D and G) Gene set enrichment analysis (GSEA) using differential expression statistics of DEGs (D) or DEPs (G) emerging from the comparison of *Rptor*-MADM/Control-MADM or cKO-*Rptor*-MADM/Control-MADM at indicated developmental time points using a curated set of GO terms. GO terms were manually grouped based on their description and the highest enrichment score (negative log10 of adjusted p-value) for each sample and GO group is shown. (E and H) Protein-protein interaction networks as determined by the STRING data base, connecting DEGs and DEPs with diverse biological functions across developmental time points. Note that networks show a higher number of interactions than expected at random for both DEGs (E, p-value = 0.0072, STRINGdb) and DEPs (H, p-value <10^-16^, STRINGdb).

**Figure S4.**
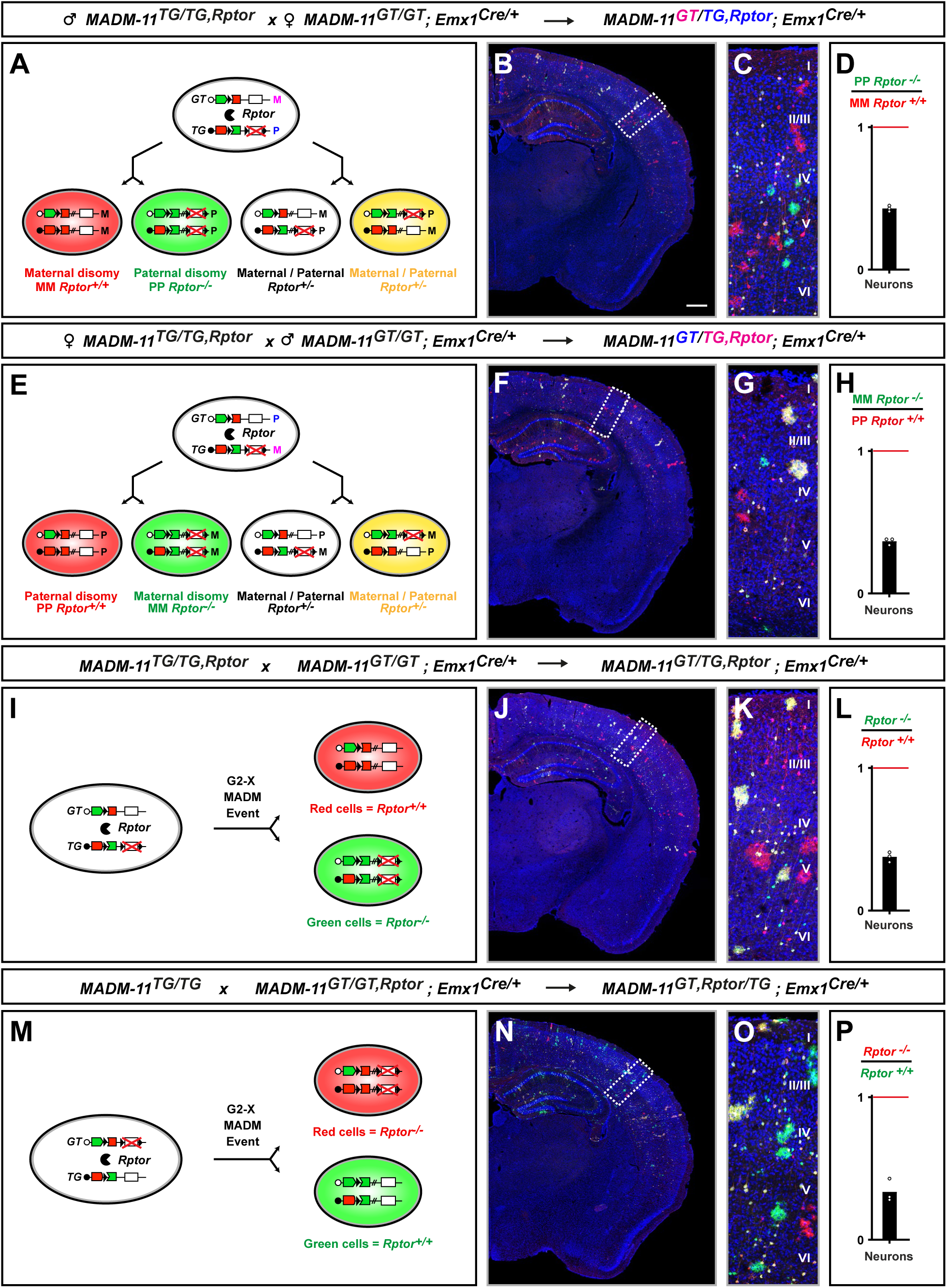
The cell-autonomous projection neuron maintenance *Rptor^-/-^* phenotype is independent of uniparental chromosome disomy and fluorescent marker, related to Figure 5. (A-H) Generation and analysis of *MADM-11^GT/TG,Rptor^*;*Emx1^Cre/+^*taking into account possible effects of uniparental chromosome disomy (UPD); i.e. parental origin of the MADM cassette with linked *Rptor-Δ* allele. (A) MADM scheme with the *Rptor-Δ* allele recombined distal to TG-MADM cassette, resulting in green *Rptor^-/-^* mutant cells and red *Rptor^+/+^* control cells. The TG-MADM cassette and linked *Rptor-Δ* allele originated from the male parent resulting in paternal UPD (PP) in the green *Rptor^-/-^* mutant cells and maternal UPD (MM) in red *Rptor^+/+^* control cells; yellow and unlabeled *Rptor^+/-^* cells resulting from G2-Z segregation were not affected. (B-D) Analysis of MADM labeling pattern in somatosensory cortex in *MADM-11^GT/TG,Rptor^*;*Emx1^Cre/+^*at P21 in overview (B) and at higher magnification (C). (D) Quantification of green PP *Rptor^-/-^* / red MM *Rptor^+/+^* ratio of excitatory projection neurons. (E-H) Generation and analysis of *MADM-11^GT/TG,Rptor^*;*Emx1^Cre/+^*taking into account the possible effects of uniparental chromosome disomy (UPD) and reversing the MADM breeding scheme. (E) MADM scheme with the *Rptor-Δ* allele recombined distal to TG-MADM cassette resulting in green *Rptor^-/-^*mutant cells and red *Rptor^+/+^* control cells. Here, the TG-MADM cassette and linked *Rptor-Δ* allele originate from the female parent resulting in maternal UPD (MM) in the green *Rptor^-/-^* mutant cells and paternal UPD (PP) in red *Rptor^+/+^* control cells; yellow and unlabeled *Rptor^+/-^* cells resulting from G2-Z segregation are not affected. (F-H) Analysis of MADM labeling pattern in somatosensory cortex in *MADM-11^GT/TG,Rptor^*;*Emx1^Cre/+^*at P21 in overview (F) and at magnification resolution (G). (H) Quantification of green MM *Rptor^-/-^* / red PP *Rptor^+/+^*ratio of excitatory projection neurons. (I-L) Generation and analysis of *MADM-11^GT/TG,Rptor^*;*Emx1^Cre/+^*. (I) Standard MADM scheme depicting G2-X event with the *Rptor-Δ* allele recombined distal to TG MADM cassette resulting in green *Rptor^-/-^* mutant cells and red *Rptor^+/+^*control cells. (J-L) Analysis of MADM labeling pattern in somatosensory cortex in *MADM-11^GT/TG,Rptor^*;*Emx1^Cre/+^* at P21 in overview (J) and at higher magnification (K). (L) Quantification of green/red ratio of excitatory projection neurons. (M-P) Generation and analysis of *MADM-11^GT,Rptor/TG^*;*Emx1^Cre/+^*. (M) MADM scheme depicting G2-X event with the *Rptor-Δ* allele recombined distal to GT MADM cassette resulting in red *Rptor^-/-^* mutant cells and green *Rptor^+/+^* control cells. (N-P) Analysis of MADM labeling pattern in somatosensory cortex in *MADM-11^GT,Rptor/TG^*;*Emx1^Cre/+^*at P21 in overview (N) and at higher magnification (O). (P) Quantification of red/green ratio of excitatory projection neurons. Cortical layers are indicated (roman numerals). Each individual data point represents one experimental animal (D, H, L and P). Bars represent mean. Significance was determined using Mann Whitney test (D, H, L and P). Scale bar, 500µm (B, F, J, and N). See also Figure S1 for complete MADM scheme, Figure S2 for breeding schemes and (Laukoter et al. 2020a; Contreras et al. 2021) for more details about MADM-based generation and labeling of UPD. Schematic is reused, adapted and modified from (Beattie et al. 2017), with permission.

**Figure S5.**
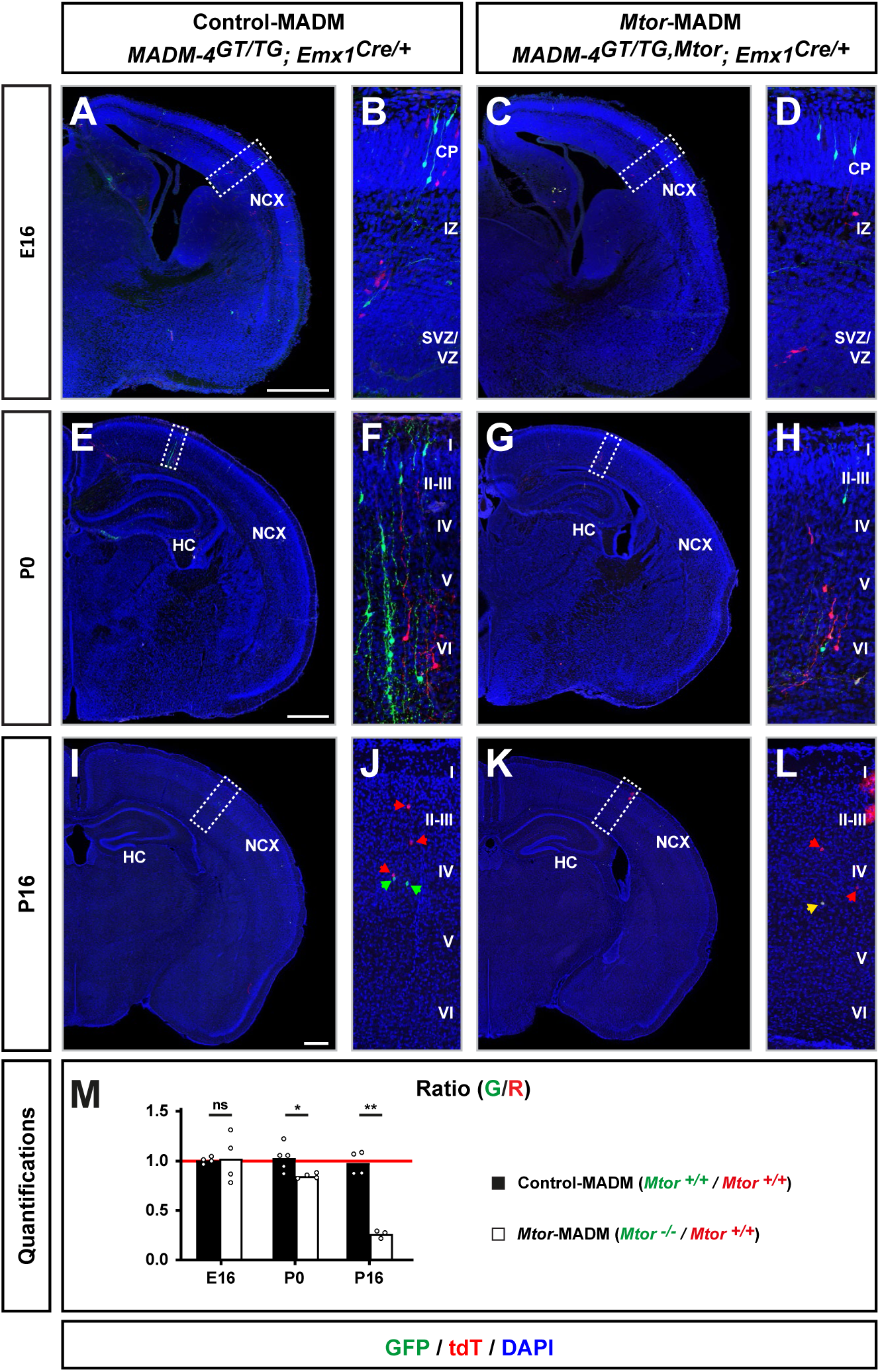
*Mtor* is cell-autonomously required for postnatal cortical projection neuron maintenance, related to Figure 5. (A-L) Analysis of MADM-labeled excitatory projection neurons in Control-MADM (A, B, E, F, I and J; *MADM-4^GT/TG^;Emx1^Cre/+^*; red cells: *Mtor^+/+^*, green cells: *Mtor^+/+^*, background: *Mtor^+/+^*) and *Mtor*-MADM (*MADM-4^GT/TG,Mtor^;Emx1^Cre/+^*(C, D, G, H, K and L), red cells: *Mtor^+/+^*, green cells: *Mtor^-/-^*, background: *Mtor^+/-^*) at E16 (A-D), P0 (E-H) and P16 (I-L). (M) Time course analysis of G/R ratio of MADM-labeled excitatory projection neurons in all cortical layers at E16, P0, and P16. Note that G/R ratio is ∼1 (equal numbers of wild-type and *Mtor^-/-^* neurons in *Mtor*-MADM) at E16 when cortical neurogenesis ends. A decrease in G/R ratio is evident only from P0 onward. Nuclei were stained using DAPI (blue). NCX: Neocortex. HC: Hippocampus. Each individual data point represents one experimental animal (M). Bars represent mean. Significance was determined using unpaired t test (M). ns: non-significant. *p < 0.05, **p < 0.01. Cortical layers are indicated in roman numerals. Scale bar, 500 µm (A, C, E, G, I, K).

**Figure S6.**
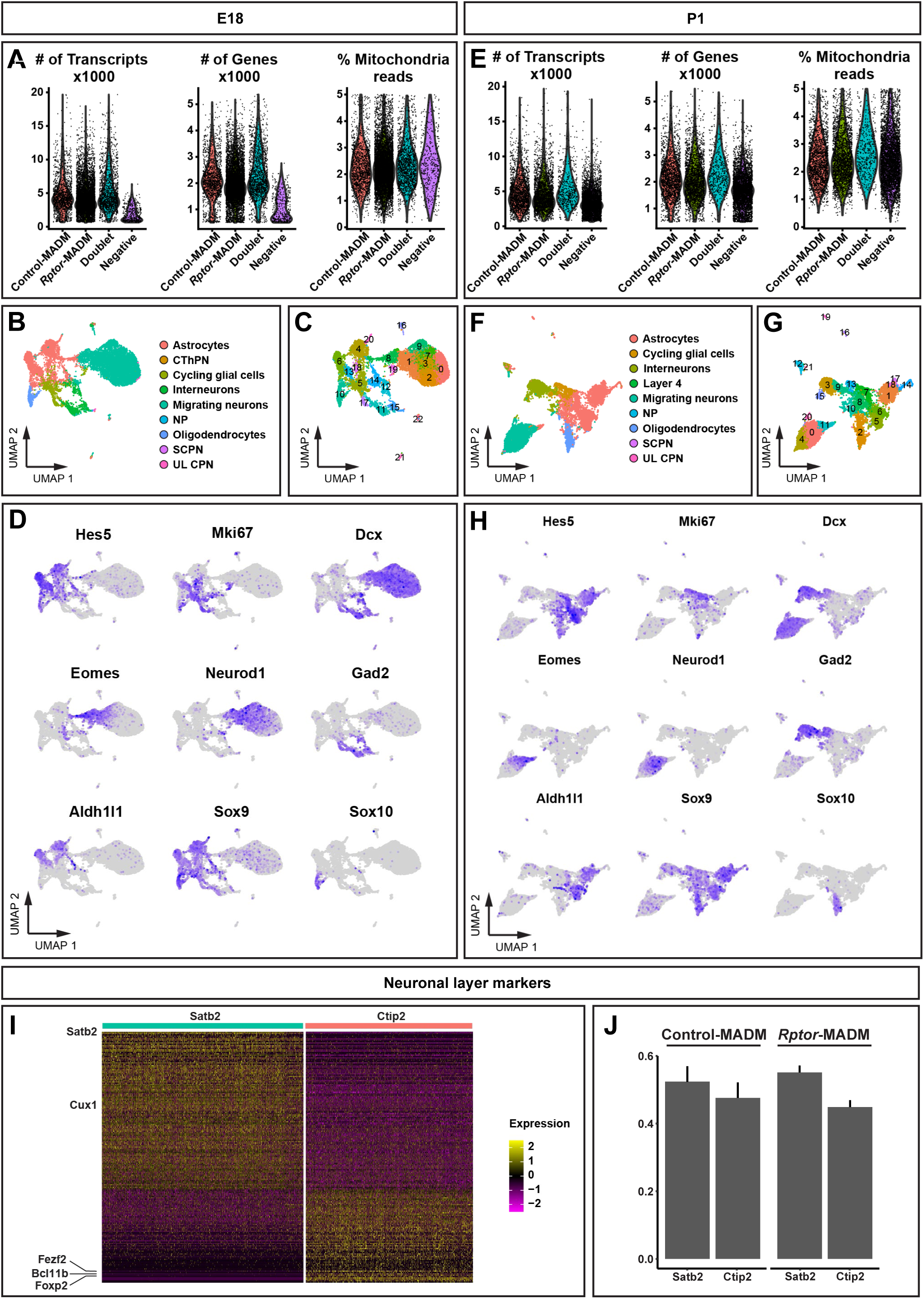
Analysis of gene expression by scRNA-seq upon *Rptor* loss of function, related to Figure 6. (A-H) Quality control metrics for cells in sequenced and demultiplexed samples (Control-MADM, *Rptor*-MADM, Doublets, Negatives) (A and B); UMAPs of sequenced cells with distinct colors indicating cell type annotation based on label transfer from (Di Bella et al. 2021) (C and E); UMAPs of sequenced cells with distinct colors and numbers indicating clusters of cells originating from unsupervised clustering (D and F); UMAPs of cells with color shading indicating expression levels of distinct marker genes for progenitors (*Hes5*), cells in active cell cycle (*Mki67*), neurons (*Dcx*), intermediate progenitors (*Eomes*), immature neurons (*Neurod1*), olfactory bulb neruoblasts (*Gad2*), glia (*Sox9*), astrocytes (*Aldh1l1*), oligodendrocytes (*Sox10*) (G and H); at E18 (A, C, D and G) and P1 (B, E, F and H). (I) Heatmap showing the expression of genes specifically expressed in *Satb2^+^* or *Ctip2^+^* projection neuron types, with indications of some well-established marker genes. (J) Relative fractions of *Satb2^+^* and *Ctip2^+^* projection neuron types in Control-MADM and *Rptor*-MADM samples. Error bars: 95% Clopper-Pearson confidence intervals.

**Figure S7.**
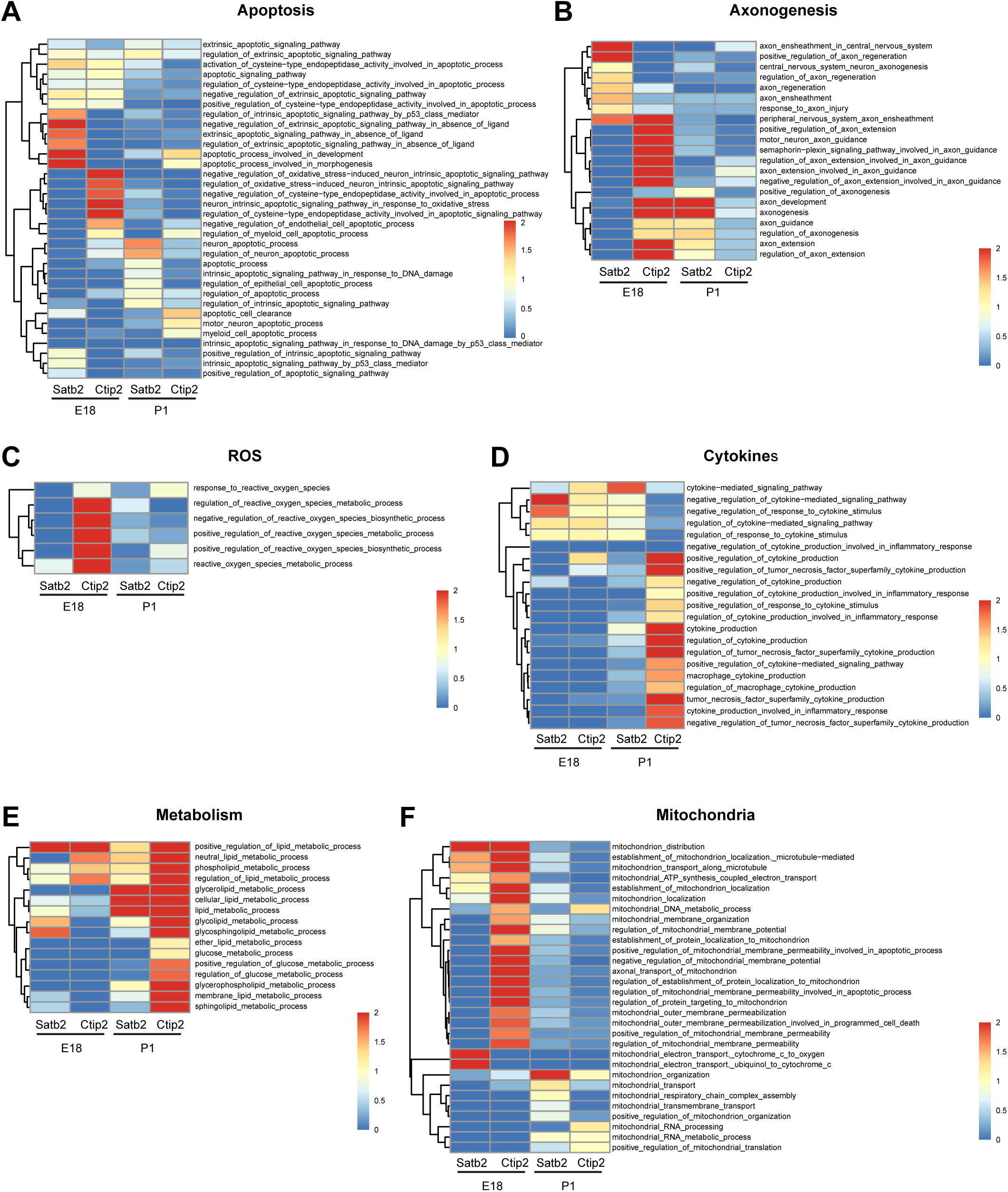
GO enrichment analysis in *Satb2^+^* and *Ctip2^+^* cortical projection neurons upon deletion of *Rptor*, related to Figure 6. (A-F) Heatmaps of GO term p-value scores as shown in Figure 6E for the entire set of investigated GO groups, complete GO term lists and with indications of GO term descriptions.

**Figure S8.**
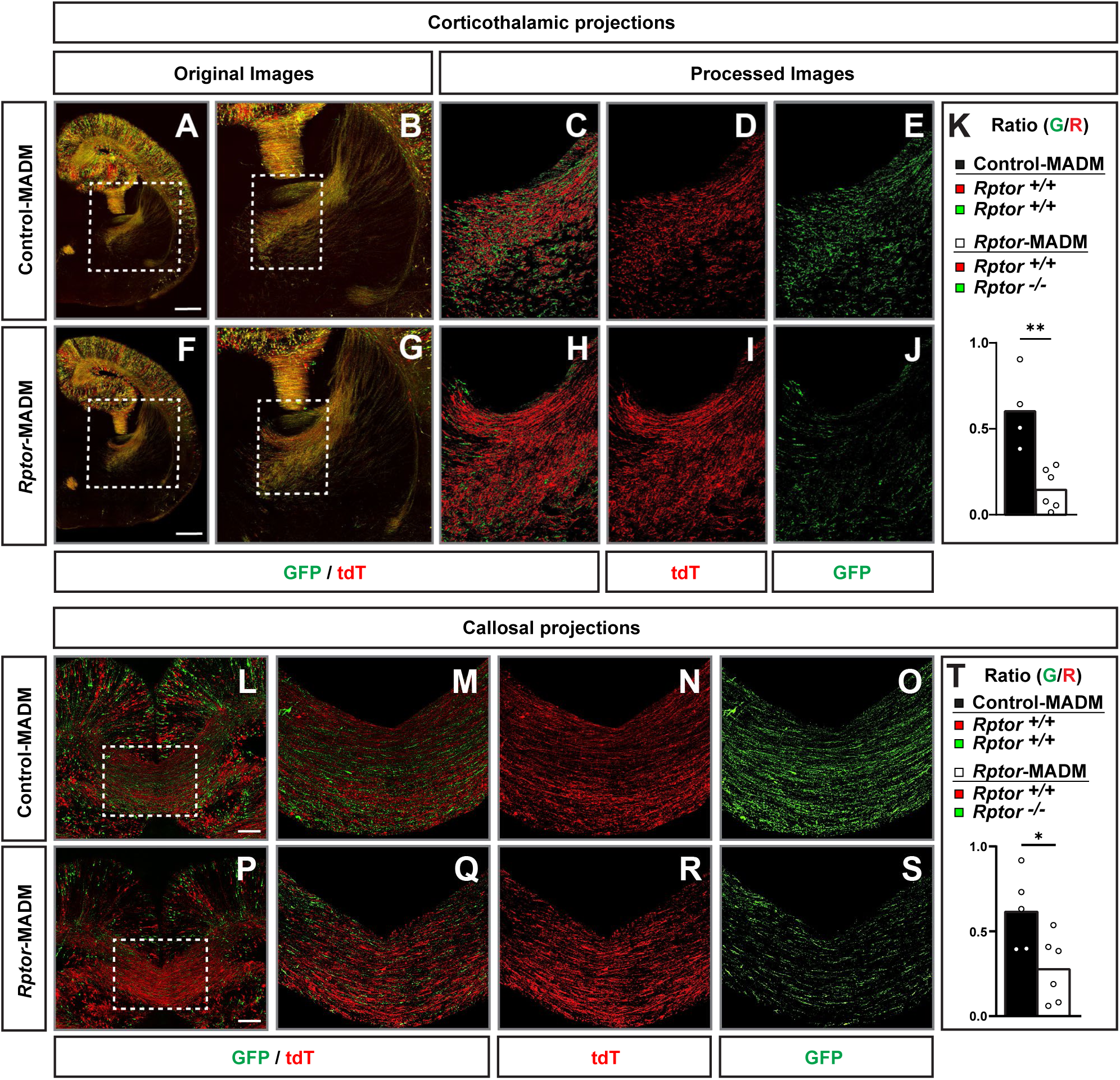
Loss of *Rptor* function results in aberrant axonal projections, related to Figure 6. (A-K) Analysis of corticothalamic axonal projections. (A, B, F, G) Representative overview images of Control-MADM (A-B; red cells: *Rptor^+/+^*; green cells: *Rptor^+/+^*; background and yellow cells: *Rptor^+/+^*) and *Rptor*-MADM (F-G; red cells: *Rptor^+/+^*; green cells: *Rptor^-/-^*; background and yellow cells: *Rptor^+/-^*) at P0. Images in (B) and (G) represent higher magnification of the boxed areas in (A) and (F), respectively. (C-E and H-J) Processed images [higher magnification of boxed areas in (B and G)] after subtracting the yellow (i.e. GFP^+^/tdT^+^ double positive) fluorescent signal, illustrating the cortico-thalamic neural projection tracks. (K) Quantification of the volume ratio of green projections compared to the red projections in Control-MADM (black bar) and *Rptor*-MADM (white bar). (L-T) Analysis of callosal axonal projections. (L-S) Processed images after subtracting the yellow fluorescent signal in Control-MADM and *Rptor*-MADM at P0. (M-O and Q-S) Higher magnification of boxed areas in (L) and (P), illustrating the callosal projection tracks. (T) Quantification of the volume ratio of green projections compared to the red projections in Control-MADM (black bar) and *Rptor*-MADM (white bar). Each individual data point represents one experimental animal in (K) and (T). Bars represent mean. Significance was determined by using unpaired t test (K and T). *p < 0.05, **p < 0.01. Scale bar, 500 μm (A and F), 100 μm (L and P).

**Figure S9.**
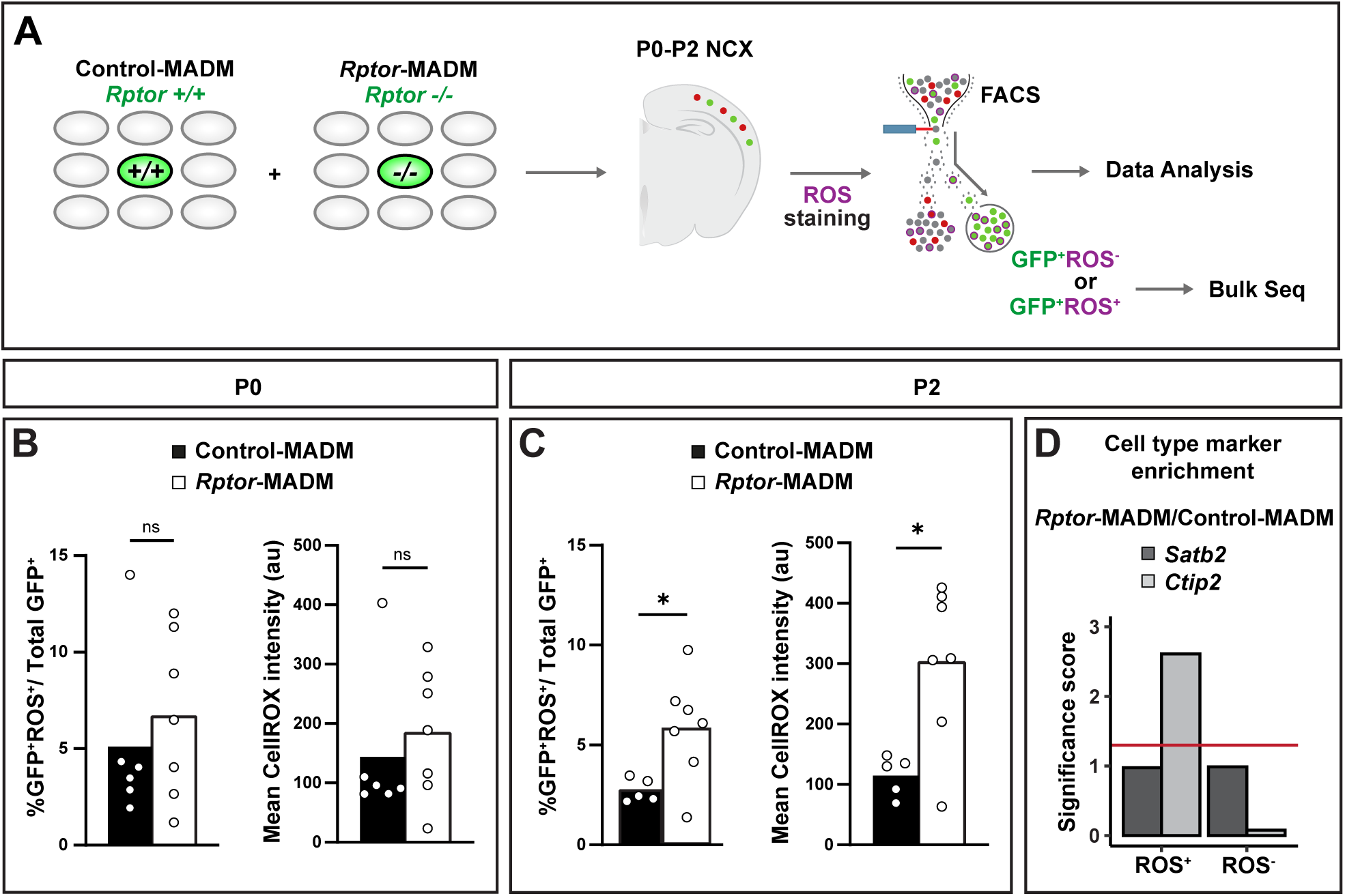
Cell-type specific increase in ROS production upon *Rptor* loss of function, related to Figure 6. (A) Illustration of the experimental strategy to FACS-enrich for *Emx1*^+^ progenitor cells and separate ROS^+^ cells from ROS^-^ cells for downstream bulk RNA-seq. (B-C) Control-MADM and *Rptor*-MADM FACS-sorted population analysis at P0 (B) and P2 (C). The plots show the percentage (%) of GFP^+^ROS^+^ cells within the total population of green (GFP^+^) MADM-labeled cells (*left*); and the mean intensity for ROS within the total population of green (GFP^+^) MADM-labeled cells (*right*). (D) Gene set enrichment analysis (GSEA) scores (negative log10 of adjusted p-value) using differential expression statistics from *Rptor*-MADM/Control-MADM comparisons in ROS^+^ and ROS^-^ samples, and marker genes of *Satb2*- and *Ctip2*-expressing neurons, respectively. Red line indicates the score corresponding to an adjusted p-value = 0.05. Scores above this threshold are considered statistically significant (adjusted p-value < 0.05). Each individual data point represents one experimental sample (B and C). Bars represent mean. Significance was determined by using one-way ANOVA with Tukey’s multiple comparison (B) or unpaired t test (C). ns: non-significant. *p < 0.05.

**Figure S10.**
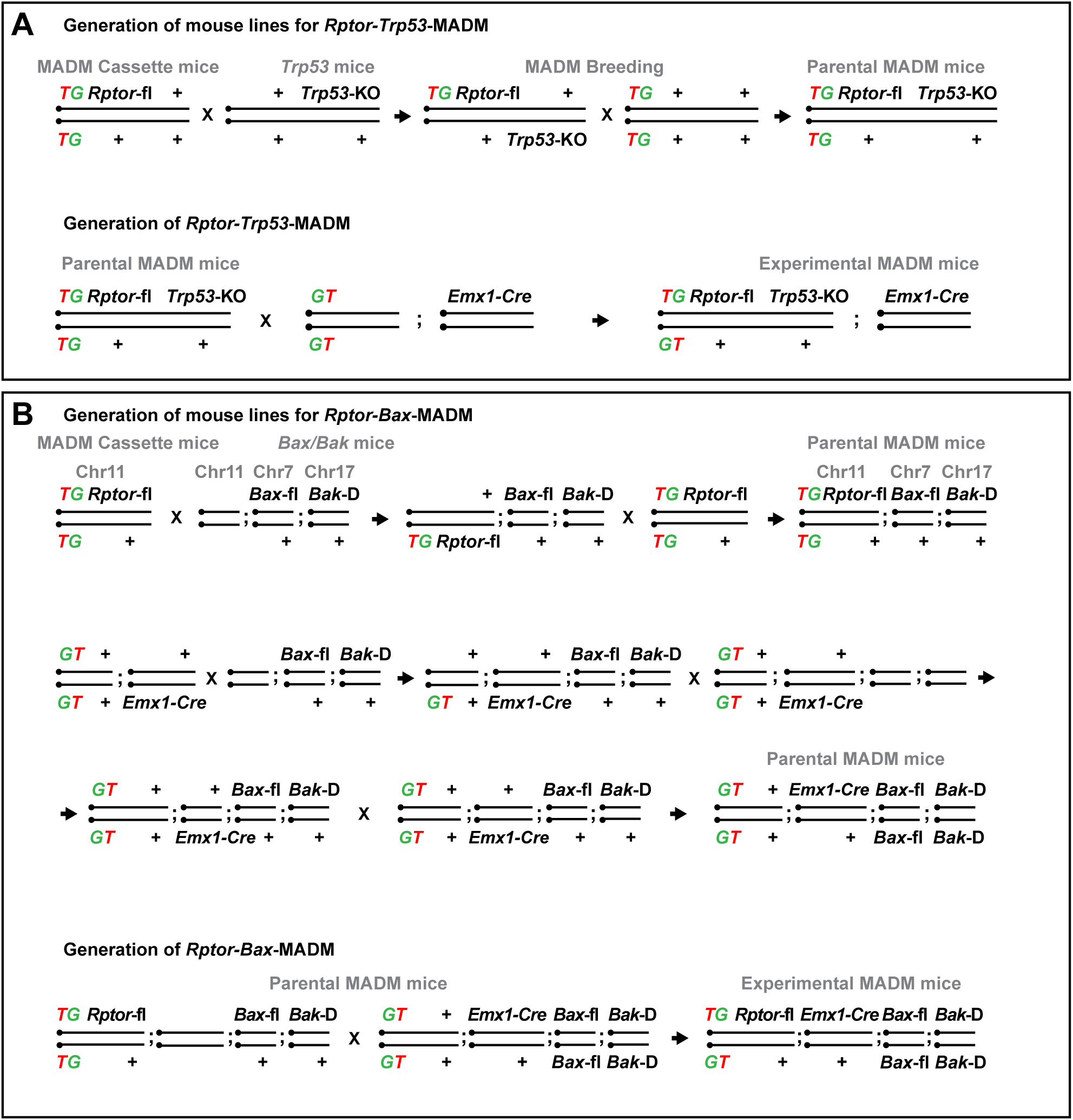
Breeding schemes to generate experimental mice for *Trp53-Rptor* and *Bax/Bak-Rptor* epistasis assays, related to Figure 7. (A-B) Breeding schemes for generating the parental mice and respective F1 experimental genotypes used for *Trp53-Rptor* [*Rptor-Trp53*-MADM (*MADM-11^GT/TG,Rptor,Trp53^*;*Emx1^Cre/+^*)](A); and *Bax/Bak-Rptor* [*Rptor*-*Bax*-MADM (*MADM-11^GT/TG^;Bax^fl/fl^;Bak^D/D^; Emx1^Cre/+^*)] (B) epistasis experiments.

